# Activity-based directed evolution of a membrane editor in mammalian cells

**DOI:** 10.1101/2022.09.26.509516

**Authors:** Reika Tei, Saket R. Bagde, J. Christopher Fromme, Jeremy M. Baskin

**Affiliations:** Department of Chemistry and Chemical Biology, Cornell University, Ithaca, NY 14853, USA; Weill Institute for Cell and Molecular Biology, Cornell University, Ithaca, NY 14853, USA; Department of Molecular Biology and Genetics, Cornell University, Ithaca, NY 14853, USA

## Abstract

Cellular membranes contain numerous lipid species, and efforts to understand the biological functions of individual lipids have been stymied by a lack of approaches for controlled modulation of membrane composition in situ. Here, we present a strategy for editing phospholipids, the most abundant lipids in biological membranes. Our membrane editor is based upon a bacterial phospholipase D (PLD), which exchanges phospholipid head groups through hydrolysis or transphosphatidylation of phosphatidylcholine with water or exogenous alcohols. Exploiting activity-dependent directed enzyme evolution in mammalian cells, we developed and structurally characterized a family of “superPLDs” with up to 100-fold higher activity than wildtype PLD. We demonstrated the utility of superPLDs for both optogenetics-enabled editing of phospholipids within specific organelle membranes in live cells and biocatalytic synthesis of natural and unnatural designer phospholipids *in vitro*. Beyond the superPLDs, activity-based directed enzyme evolution in mammalian cells is a generalizable approach to engineer additional chemoenzymatic biomolecule editors.

## INTRODUCTION

Cellular membranes have myriad functions, ranging from being selectively permeable barriers to platforms for initiating signaling pathways^1,2^. Though membranes contain hydrophobic proteins and glycoconjugates^3^, by far their most abundant constituents are lipids. Understanding how each individual lipid constituent contributes to specific properties and functions of membranes remains a major challenge in membrane biology, one that requires tools for altering the lipid content of endogenous membranes with high molecular and spatiotemporal precision^4^. Akin to how single amino acid substitutions via site-directed mutagenesis or amber suppression have transformed our ability to perform structure-function relationships within the proteome^5,6^, controllable lipid-modifying enzymes can serve as “membrane editors” to enable the selective manipulation of individual lipid species within membranes^7,8^.

This strategy has seen the most success with the phosphoinositides, a family of phosphorylated derivatives of phosphatidylinositol, where chemical- or light-induced proximity has been harnessed to create a suite of tools for rapid phosphorylation/dephosphorylation of the inositol head group in situ^9–13^. Though critically important for many signaling pathways, phosphoinositides are rare lipids, and similar tools for membrane editing beyond this tiny sector of the lipidome are scant^4^. Phosphatidylcholine (PC) is the most abundant lipid within cellular membranes, and we envisioned that it could serve as a substrate for a general membrane editor capable of replacing the choline head group with natural and unnatural head groups to create a wide array of desired phospholipids on demand.

Phospholipase D (PLD) is an ideal starting point for such a general phospholipid membrane editor. PLD catalyzes hydrolysis of PC to form a signaling lipid, phosphatidic acid (PA)^14,15^, and it can also catalyze transphosphatidylation with exogenous alcohols to swap out head groups to form a variety of natural and unnatural phospholipids^16–19^. Though mammalian cells have endogenous PLD enzymes, they are dispensable for viability and exhibit low levels of basal activity^16^. We have previously identified a microbial PLD that possesses hydrolysis and transphosphatidylation activities in mammalian cells and is amenable to light-mediated control of its localization and activity^20^. However, the activity of this PLD in mammalian cells is modest, owing to its acidic pH maximum and multiple disulfide bonds^21,22^, limiting its utility to circumstances where very low levels of PA formed by hydrolysis are sufficient to induce a signaling outcome. Thus, we envisioned applying directed evolution to develop super-active PLDs (superPLDs) that would be highly efficient membrane editors for PC hydrolysis to PA and transphosphatidylation to other phospholipids.

Directed enzyme evolution is typically performed as an iteration of two basic steps that mimic natural selection: random or targeted mutagenesis of a gene to generate a library of variants and identification of rare variants that exhibit a desired function through selection or screening^23,24^. Where feasible, selection is preferred to screening, to increase throughput^25^. In vivo selections are typically performed using *E. coli* or *S. cerevisiae* as host cells, even when the evolved enzyme is ultimately intended for use in mammalian cells. However, this approach can be problematic for lipid-modifying enzymes, as their substrates are components of membranes, whose compositions and properties differ substantially between bacteria, fungi, and higher eukaryotes^26,27^. In these cases, directed evolution in mammalian cells, despite the extra challenges that it brings, is warranted^28^.

Although activity-based labeling is commonly used for directed enzyme evolution, it is typically implemented either *in vitro* or with cell surface display in bacterial or yeast cells^23,29^. Here, we overcome substantial technical challenges to develop a directed enzyme evolution strategy for PLD in mammalian cells that harnesses a bioorthogonal, activity-based imaging method for fluorescently tagging cellular membranes proportional to PLD activity. Using this platform, we obtained a series of superPLDs with greatly enhanced stability in the intracellular environment and catalytic efficiencies of up to 100-fold higher than wildtype PLD. Structural and biochemical analysis revealed that superPLDs possess an expanded active site that allows greater access of water and alcohol substrates and are less reliant upon intramolecular disulfides. Because of their significantly improved intracellular stability and catalytic efficiencies, superPLDs open up applications both in cell biology for membrane editing of the phospholipidome and in biotechnology for the biocatalytic production of commodity and designer phospholipids. Moreover, our demonstration of activity-based directed enzyme evolution in mammalian cells sets the stage for engineering of other chemoenzymatic labeling systems in mammalian cells.

## RESULTS

### Activity-based directed evolution in mammalian cells achieves a substantial enhancement of PLD activity

We have previously identified PLD from *Streptomyces sp. PMF* for heterologous expression in mammalian cells and developed a light-controlled, optogenetic version of it (optoPLD; Supplementary Fig. 1)^20^. Though optoPLD enabled production of PA or certain phosphatidyl alcohol lipids with organelle-level precision, it exhibited very modest activity, compromising its temporal resolution, and it only accepted a limited set of alcohol substrates. To improve its activity, we first used PLD-expressing yeast cells for directed evolution^20^. However, the FACS-based selection was inefficient due the yeast cell wall, which prevented efficient entry and rinse-out of labeling reagents into live and even fixed cells. Consequently, this platform did not yield PLD mutants with substantially enhanced activity compared to PLD^WT^; the best-performing mutant, G429D, was only 1.3-fold better than PLD^WT^. Therefore, we turned to directed evolution in mammalian cells.

Our activity-based directed evolution strategy in mammalian cells comprises four fundamental steps (Fig. 1). First, a PLD library with random mutations is generated by error-prone PCR, using either WT or G429D as a template, and then cloned into a lentiviral vector containing the optoPLD system, which enables blue light-dependent PLD activation mediated by CRY2-CIBN dimerization^30^ to recruit PLD to a desired membrane (Supplementary Fig. 1). Second, lentivirus containing this optoPLD library is generated and delivered into HEK 293T cells such that each cell expresses a different optoPLD mutant. Third, activity-based fluorescent labeling is performed using a bioorthogonal labeling method termed Imaging PLD Activity with Clickable Alcohols via Transphosphatidylation (IMPACT)^31,32^ (Fig. 2a). Fourth, cells are sorted by FACS based on IMPACT fluorescence intensity normalized to optoPLD expression, with IMPACT-high cells collected and propagated.

**Figure 1.**
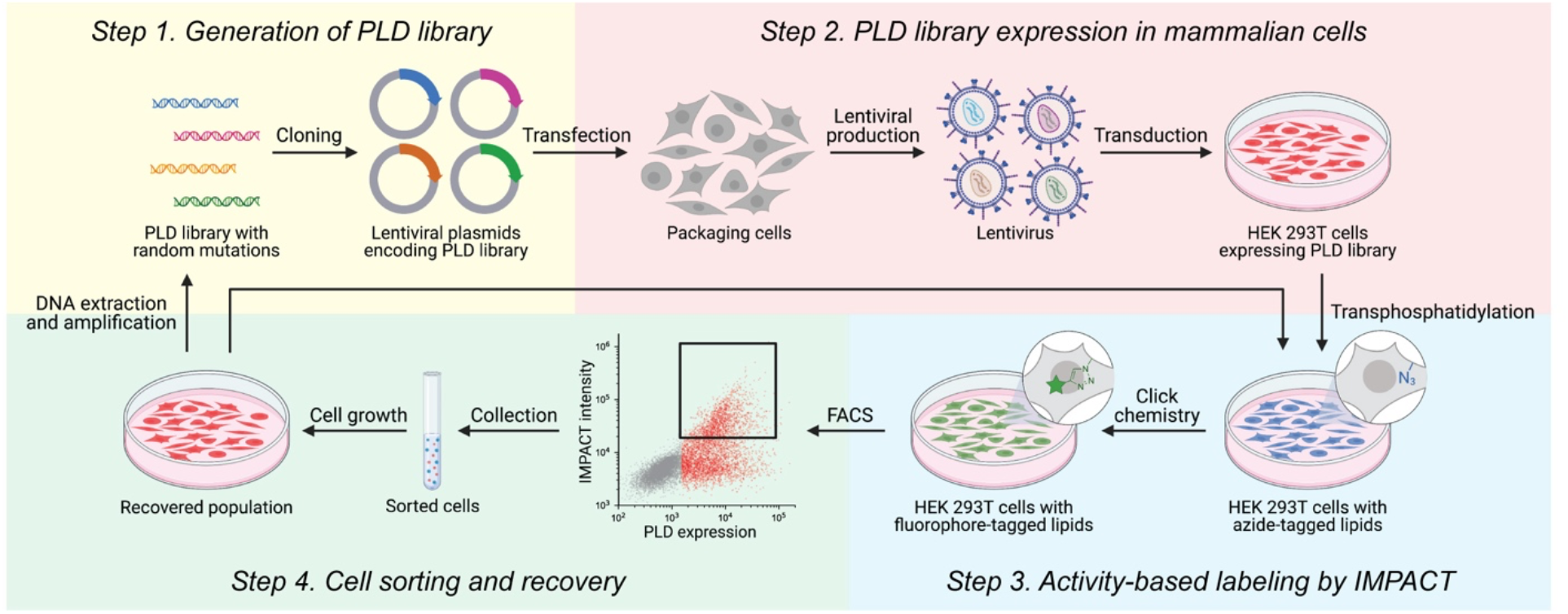
Design of an activity-dependent directed enzyme evolution strategy to create a phospholipase D (PLD)-based membrane editor in mammalian cells. Step 1: PLDs with random mutations are generated by error-prone PCR and inserted into a lentiviral optoPLD plasmid. Step 2: Packaging cells are transfected with the lentiviral plasmids to produce lentivirus encoding the optoPLD library, which is then transduced into HEK 293T cells. Step 3: Cells expressing the optoPLD library are labeled with IMPACT to fluorescently tag cellular membranes based on the catalytic activity of PLD. Step 4: Cells with high IMPACT labeling intensity are isolated by FACS, expanded, and then either the labeling is repeated to better enrich IMPACT-high cells or DNA is extracted and amplified for further rounds of evolution or clonal isolation and sequencing.

**Figure 2.**
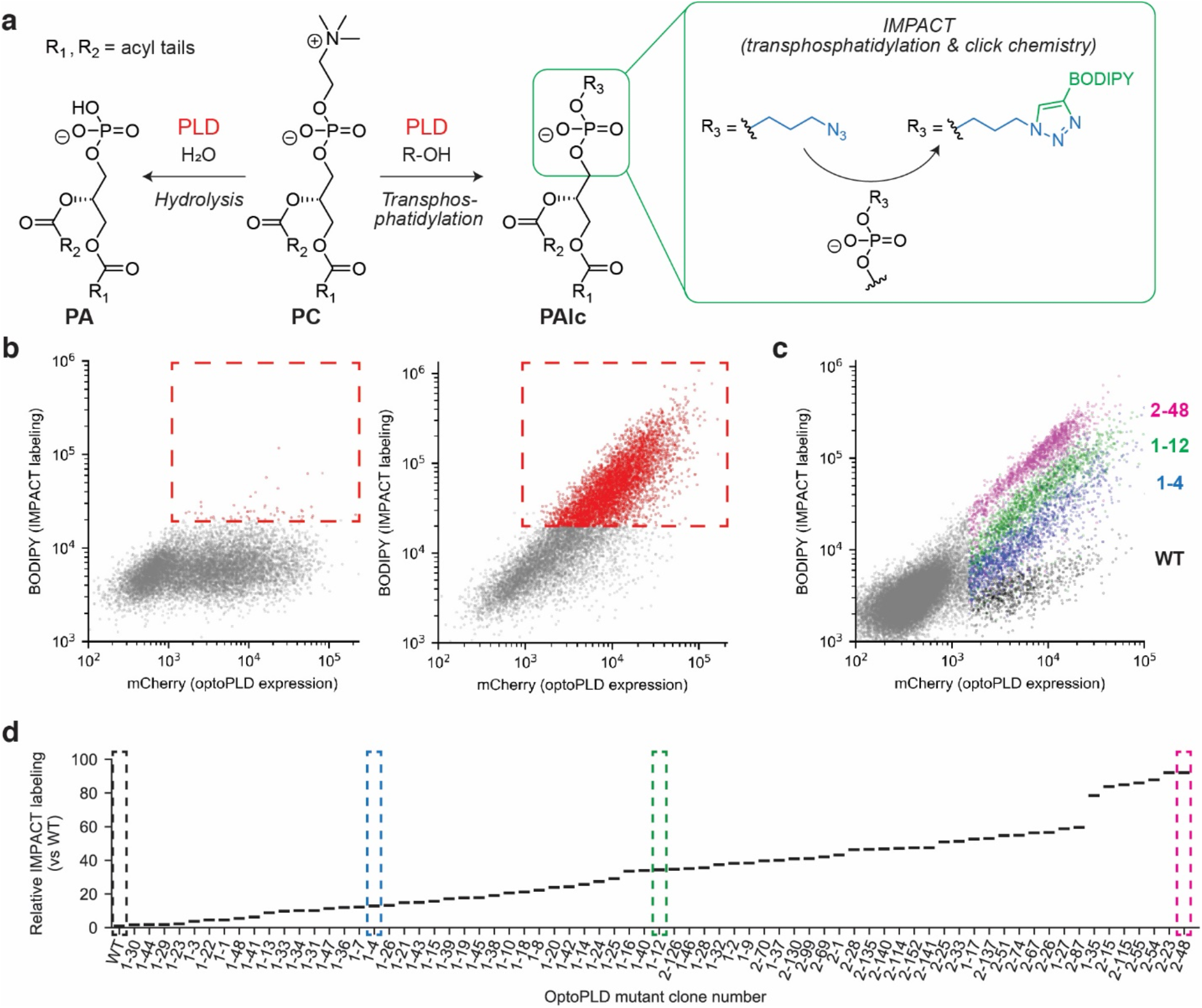
Directed evolution yields PLD mutants with greatly enhanced activities. **a**, Reactions catalyzed by PLD, including hydrolysis (left) and transphosphatidylation (right) by using water and alcohols as substrates, respectively. For IMPACT, transphosphatidylation is used to produce azide-tagged phospholipids, which are then reacted with a bicyclononyne-BODIPY probe to fluorescently tag the lipids produced by PLD. **b**, FACS plots showing signal enrichment before (left) and after (right) five rounds of selection without mutagenesis. Red dots indicate cells that were collected. **c**, Overlay of FACS plots of cells expressing different optoPLD mutants. Colored dots indicate cells expressing PLD^WT^ (black), 1-4 (blue), 1-12 (green) and 2-48 (magenta). **d**, Relative PLD activity of representative plasma membrane-targeted optoPLD mutants obtained by directed evolution. IMPACT fluorescence intensity normalized to optoPLD expression was determined by flow cytometry, and the activities are plotted as relative IMPACT labeling of the indicated PLD mutant compared to PLD^WT^.

After 1–2 cycles of IMPACT and FACS sorting, the enriched PLD library is isolated by DNA extraction and PCR amplification. Subsequent rounds of evolution, with or without additional mutagenesis, is performed while increasing stringency by lowering the concentration of the IMPACT labeling reagent azidopropanol (Supplementary Fig. 2a–s). Overall, we performed three rounds of selection with mutagenesis to increase library diversity and, critically, then five rounds without additional mutagenesis to remove false-positive populations, leading to a dramatic enrichment of highly active PLD mutants (Fig. 2b).

After the final round of selection, we isolated 216 individual clones of the enriched library (in two batches) and found that 194 (90%) exhibited higher activity than PLD^WT^ using IMPACT labeling to assess activity of these mutant PLDs within an optoPLD system. These mutants exhibited a wide range of activities, with the highest active mutant, clone 2-48, exhibiting ~100x higher activity than PLD^WT^ (Fig. 2c–d and Supplementary Fig. 2t–v). Due to this substantial improvement in performance in cells, we denoted these mutants as superPLDs.

### SuperPLD is an efficient PC hydrolase and transphosphatidylase

Having obtained PLD mutants with a wide-ranging degree of transphosphatidylation activities to generate fluorescent lipids via IMPACT, we next assessed their ability to catalyze PC hydrolysis to form PA and transphosphatidylation to form other useful phospholipids (Fig. 2a). To test superPLD-mediated PA production in cells, we used a PA-binding probe, GFP-PASS^33^, to visualize the subcellular localizations of PA. An optoPLD construct containing the highest active superPLD (2-48) exhibited substantial light-independent background activity in cells (Supplementary Fig. 2w–x), resulting in increased cytotoxicity under certain conditions, e.g., stable expression following lentiviral transduction. Therefore, we generated optoPLDs bearing moderately active superPLDs (superPLD^x10^ and superPLD^x30^, made from 1-4 and 1-12 with approximately 10x and 30x higher activity than PLD^WT^, respectively) capable of being recruited to either the plasma membrane or lysosomes upon light activation (Fig. 3a). LC–MS analysis of transphosphatidylation reactions revealed that superPLD^x30^ activation consumed approximately 2–4% of PC in cells, and total PC levels did not significantly change, likely due to continuous replenishment from biosynthesis (Supplementary Fig. 3a–b). With plasma membrane-targeted optoPLDs, superPLD^x10^ and superPLD^x30^ but not PLD^WT^ efficiently recruited GFP-PASS to the plasma membrane (Fig. 3b and Supplementary Fig. 3c). Similarly, superPLD^x10^ and superPLD^x30^ greatly outperformed PLD^WT^ in lysosome-targeted optoPLD constructs (Fig. 3c and Supplementary Fig. 3d–e).

**Figure 3.**
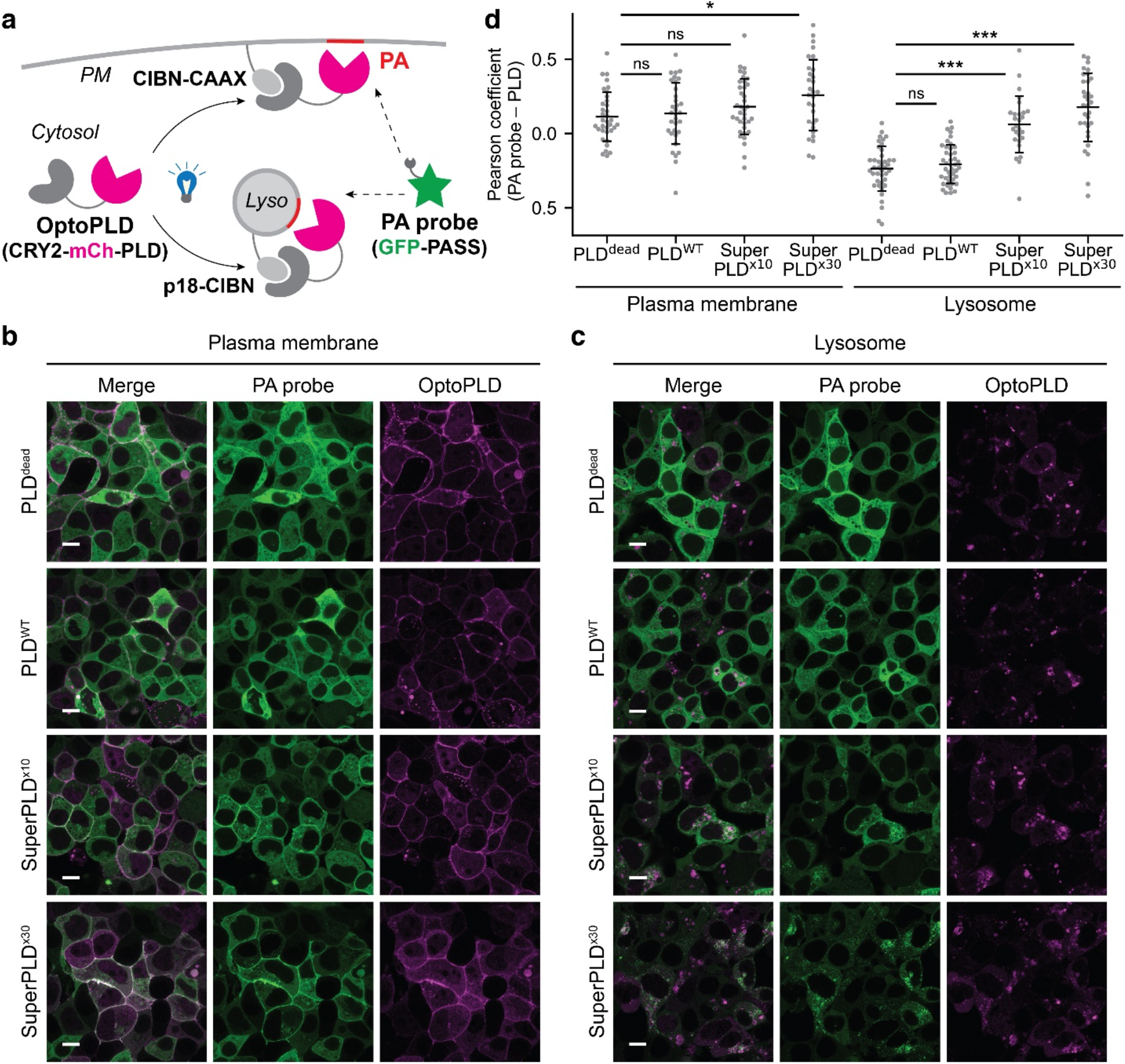
Evaluation of superPLD activity in cells. **a**, Schematic depicting the experimental design of targeting optoPLD to the plasma membrane (PM) or lysosomes (lyso) and using the PA probe GFP-PASS to visualize PA produced by optoPLD. **b–c**, Confocal images of HEK 293T cells co-expressing the PA probe and optoPLD targeted to the plasma membrane (**b**) or lysosomes (**c**) after 30 min incubation with 488 nm light. **d**, Quantification of co-localization between PA probe and optoPLD. The plots show Pearson’s correlation coefficient of PA probe and optoPLD. Black horizontal lines indicate mean and vertical error lines indicate standard deviation (n=30–40). PLD^dead^; a catalytically dead PLD bearing the H167A mutation, PLD^WT^; wild-type PLD, superPLD^x10^; superPLD mutant clone 1-4, superPLD^x30^; superPLD mutant clone 1-12. Scale bar: 10 μm.

To evaluate PLD activity *in vitro*, we purified His-tagged superPLDs expressed in the *E. coli Rosetta 2* strain (Supplementary Fig. 4a). Notably, PLD^WT^ was unable to be purified in this manner until switching to *E. coli Rosetta-gami 2*, an engineered strain that facilitates disulfide bond formation in the cytosol^34^, consistent with this PLD being a secreted protein with four disulfide bonds (Supplementary Fig. 4b–c)^16,35^. That superPLD can be robustly purified from a conventional *E. coli* strain suggests that one potential source of improved superPLD performance is an enhanced stability in intracellular, reductive environments. Supporting this hypothesis, the activity increase of superPLD was more substantial in cells than *in vitro*, and superPLD exhibited enhanced chemical stability compared to PLD^WT^ (Fig. 2d and Supplementary Fig. 4d–h).

Given that traditionally, transphosphatidylation by PLDs is preferred to hydrolysis^22,36^, we evaluated the ability of superPLD as a catalyst for the *in vitro* synthesis of a variety of phospholipids from phosphatidylcholine and alcohol substrates. By quantifying both PA and phosphatidyl alcohol products by LC–MS, we found that superPLD could be successfully used for synthesis of various useful natural and unnatural phosphatidyl alcohols derived from both primary and secondary alcohols, with minimal PA by-products formed (Fig. 4). Notably, several unnatural phospholipids with reactive handles such as azide^31^ and alkyne^37^ could be synthesized with high selectivity and yield. These studies demonstrate the utility of superPLD as a biocatalyst to efficiently produce natural and unnatural phospholipids in *in vitro* chemoenzymatic reactions.

**Figure 4.**
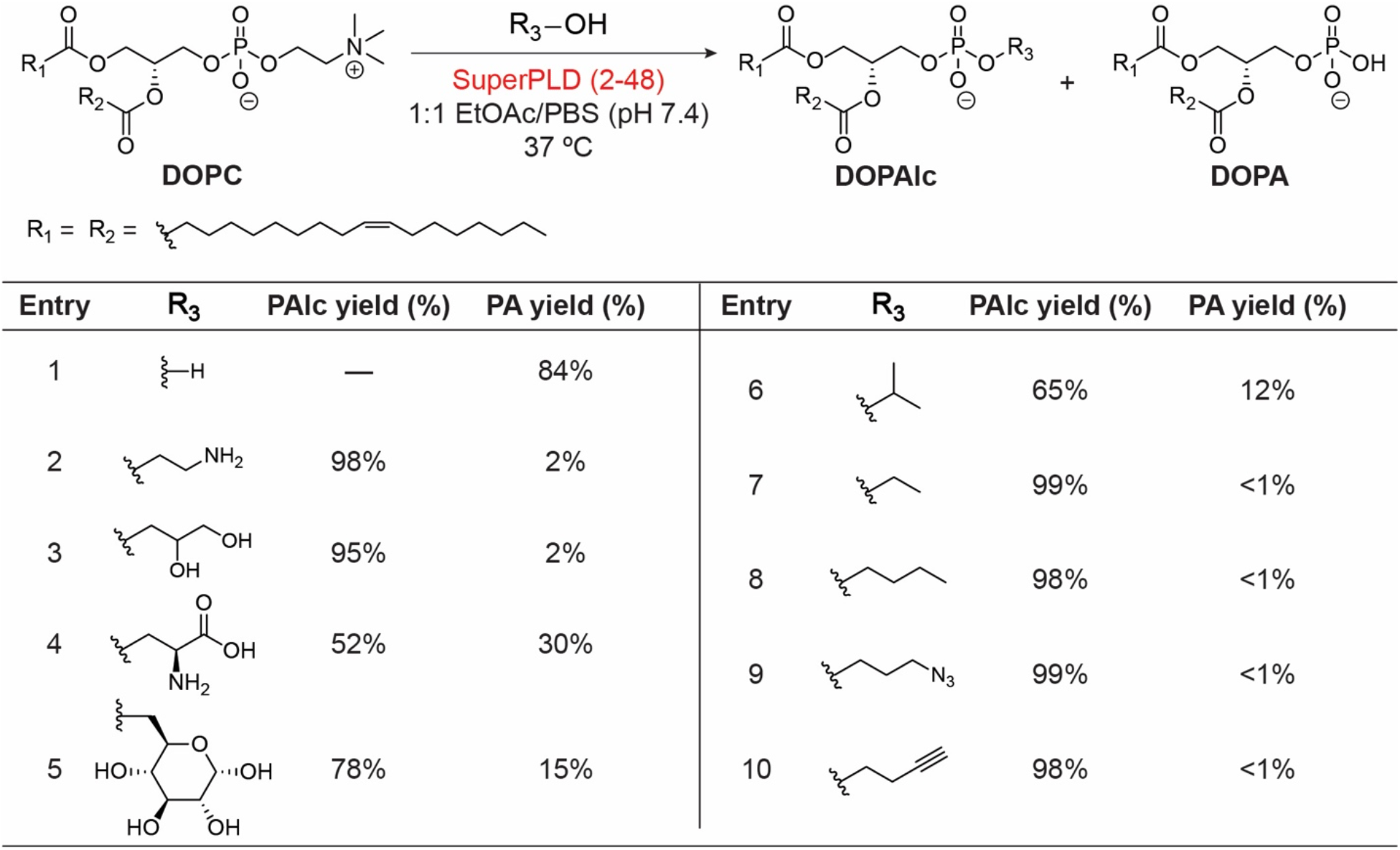
Synthesis of designer phospholipids by *in vitro* reaction. The reactions were performed in a biphasic system of ethyl acetate (EtOAc, 0.8 mL) and phosphate-buffered saline (PBS, pH 7.4, 1.0 mL) with 0.8 mg dioleoyl phosphatidylcholine (DOPC), 0.1 μg superPLD (2-48) and 200 μmol (for entries 1–6) or 50 μmol (for entries 7–10) of alcohol. PAlc yield and PA yield indicate the % conversion of DOPC to dioleoyl phosphatidyl alcohol (DOPAlc) and dioleoyl phosphatidic acid (DOPA), respectively.

### SuperPLD modulates PA-dependent signaling pathways

To establish the general ability of superPLDs to generate physiologically active PA pools, we assessed optogenetic versions of superPLD for modulation of three different PA-dependent signaling pathways. First, we confirmed that PA made at the plasma membrane by superPLD^x30^ can attenuate Hippo growth restriction pathway by triggering translocation of Yes-associated protein (YAP) from the cytosol to the nucleus in serum-starved cells (Supplementary Fig. 5a–b), as we had previously shown for the WT form of optoPLD^20^.

**Figure 5.**
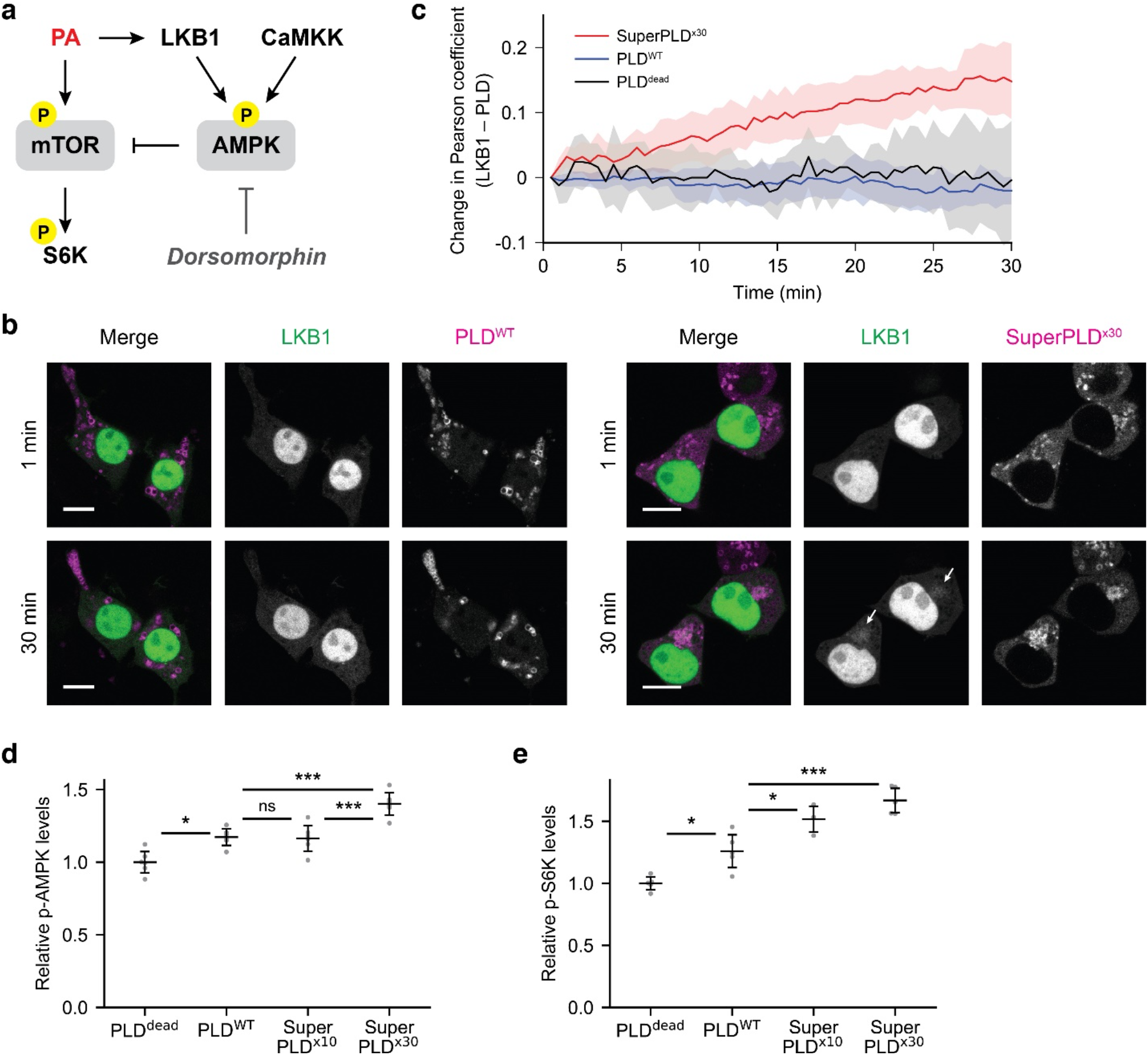
Application of superPLD to manipulate PA signaling. **a**, Schematic diagram of effects of PA on AMPK and mTOR signaling. **b–c**, Recruitment of GFP-LKB1 triggered by acute PA production on lysosomes by superPLD. GFP-LKB1 and optoPLD (mCherry) fluorescence were measured for 30 min, and changes in Pearson correlation coefficient, representing recruitment of GFP-LKB1 to optoPLD-positive membranes, are plotted for each condition (n=5). Scale bar: 10 μm. **d**, Quantification of phospho-AMPK (p-AMPK) levels in HEK 293T cells with plasma membrane-targeted optoPLDs. Cells were pretreated with a CaMKK inhibitor (STO-609) for 6 h to eliminate CaMKK-mediated AMPK activation, followed by 30 min incubation with intermittent 488 nm light. p-AMPK levels were determined by Western blot (n=6). **e**, Quantification of phospho-S6 kinase (p-S6K) levels in HEK 293T cells with plasma membrane-targeted optoPLDs. Cells were pretreated with dorsomorphin for 1 h to eliminate effects of p-AMPK on mTOR activity, followed by 30 min incubation with intermittent 488 light. p-S6K levels were determined by Western blot (n=6). See Supplementary Fig. 5c–d for representative Western blots.

In addition to Hippo signaling, PA can also regulate two additional pathways related to nutrient sensing and cell growth: mammalian target of rapamycin (mTOR) signaling^38,39^ and AMP-activated protein kinase (AMPK) signaling^40,41^. PA binds to liver kinase B1 (LKB1), which phosphorylates AMPK to activate it; however, such effects counteract the direct stimulatory effect of PA on mTOR signaling due to crosstalk between AMPK and mTOR signaling (Fig. 5a)^41^. To verify this finding and assess the ability of an optoPLDs to induce LKB1 translocation to PA-rich membranes, we co-expressed GFP-tagged LKB1 and optoPLDs in HEK 293T cells. Strikingly, PA production on lysosomes by superPLD, but not by PLD^WT^, was sufficient to trigger LKB1 recruitment to lysosomes, consistent with the reported LKB1–PA interaction (Fig. 5b–c).

We then transduced HEK 293T cells with optoPLD using lentivirus and analyzed effects on PA signaling by Western blotting, in these instances targeting optoPLD to the plasma membrane to mimic the localization of endogenous PLDs when they are stimulated^16,18^. Although as expected, the cells had basal levels of AMPK phosphorylation^42,43^, PA production by superPLD led to a significant increase in p-AMPK (Fig. 5d and Supplementary Fig. 5c). Moreover, under conditions when AMPK signaling was inhibited with dorsomorphin to reduce crosstalk with mTOR signaling, PM-targeted superPLD increased phosphorylation of the mTOR effector S6 kinase (Fig. 5e and Supplementary Fig. 5d). These results demonstrate that PA made by optoPLDs can independently activate mTOR and AMPK signaling, and importantly, the superPLD elicited a much stronger response in both cases. Collectively, these studies establish optogenetic superPLDs as generally useful tools for acute manipulation of mammalian PA-dependent signaling pathways.

### Several mutations cooperate to enhance superPLD activity

We next sought to characterize the mutations that led to increased PLD activity in the superPLDs. Sequencing analysis revealed that each superPLD clone carried 6–10 mutations (Supplementary Fig. 6a). In addition to the G429D mutation present in one of the two templates, the selection produced a few other mutations present in many superPLDs (A258T, G381V, and T450A). Generation of mutant PLDs containing some or all of these four mutations only produced modest improvements, in a multiplicative manner, over PLD^WT^, suggesting likely founder effects for these common mutations (Supplementary Fig. 6b).

To dissect the effects of a broader set of mutations, we constructed individual point mutants in two backgrounds, WT and G381V, which was present in all superPLD clones, and assayed their activity (Supplementary Fig. 7a). Individual mutations had variable effects on PLD activity, ranging from a slight decrease to an up to ~7-fold increase, with no magic bullet mutation of recapitulating the activity of the best superPLDs. A minority of mutations did not follow a multiplicative pattern, e.g., G406S, which did not increase PLD activity alone but caused a 3-fold increase in mutant backgrounds (Supplementary Fig. 7a–c). We classified mutations into three groups: (1) mutations that consistently increased PLD activity in various backgrounds, (2) mutations that consistently decreased PLD activity or had negligible effects, and (3) mutations that exhibited divergent effects on activity alone vs. in mutant backgrounds (Supplementary Fig. 7d). Collectively, this mutational analysis established that the strong effects on superPLD activity are due to the cooperative effect of multiple mutations, most of which are far from the enzyme active site, based on the crystal structure of PLD^WT 44^.

### X-ray analysis of superPLD reveals an expanded active site

To investigate how the 3D structure of superPLD might affect its mechanism, we performed X-ray crystallography on the two most active mutants, 2-23 and 2-48, with structures determined at resolutions of 1.91 and 1.85 Å, respectively (Fig. 6 and Supplementary Fig. 8a–b). Despite sharing only five out of nine mutations, these two superPLDs had structures that were nearly identical to one another but, in certain key ways, different from the reported PLD^WT^ structure^44,45^. Though overall, the three structures were largely similar, there were major changes in four flexible loops that form the entry of the active site (Supplementary Fig. 8). In the superPLD structures, two flexible loops near the lipid head group binding site (loops 1 and 4) were farther away from the active site, causing a wider opening than in PLD^WT^. However, the two loops closer to the lipid acyl tail binding site (loops 2 and 3) were slightly closer to the active site compared to their positions in PLD^WT^. Critically, the positions of three bulky aromatic residues within loops 1 and 4 (W186, Y190 and Y383) were substantially shifted in the superPLD structures, creating extra space in the catalytic pocket (Fig. 6a–e; note that Y383 is not resolved but its movement is inferred by examining the position of H440, which occupies the position occupied by Y383 in the PLD^WT^ structure). Overall, we consider that the changes to loops 1–4, which delineate the active site opening at the enzyme–membrane interface, are an important factor that leads to an expanded catalytic pocket where the lipid head group binds, enabling greater access of water and alcohols to the active site and contributing to the increased activity of superPLDs.

**Figure 6.**
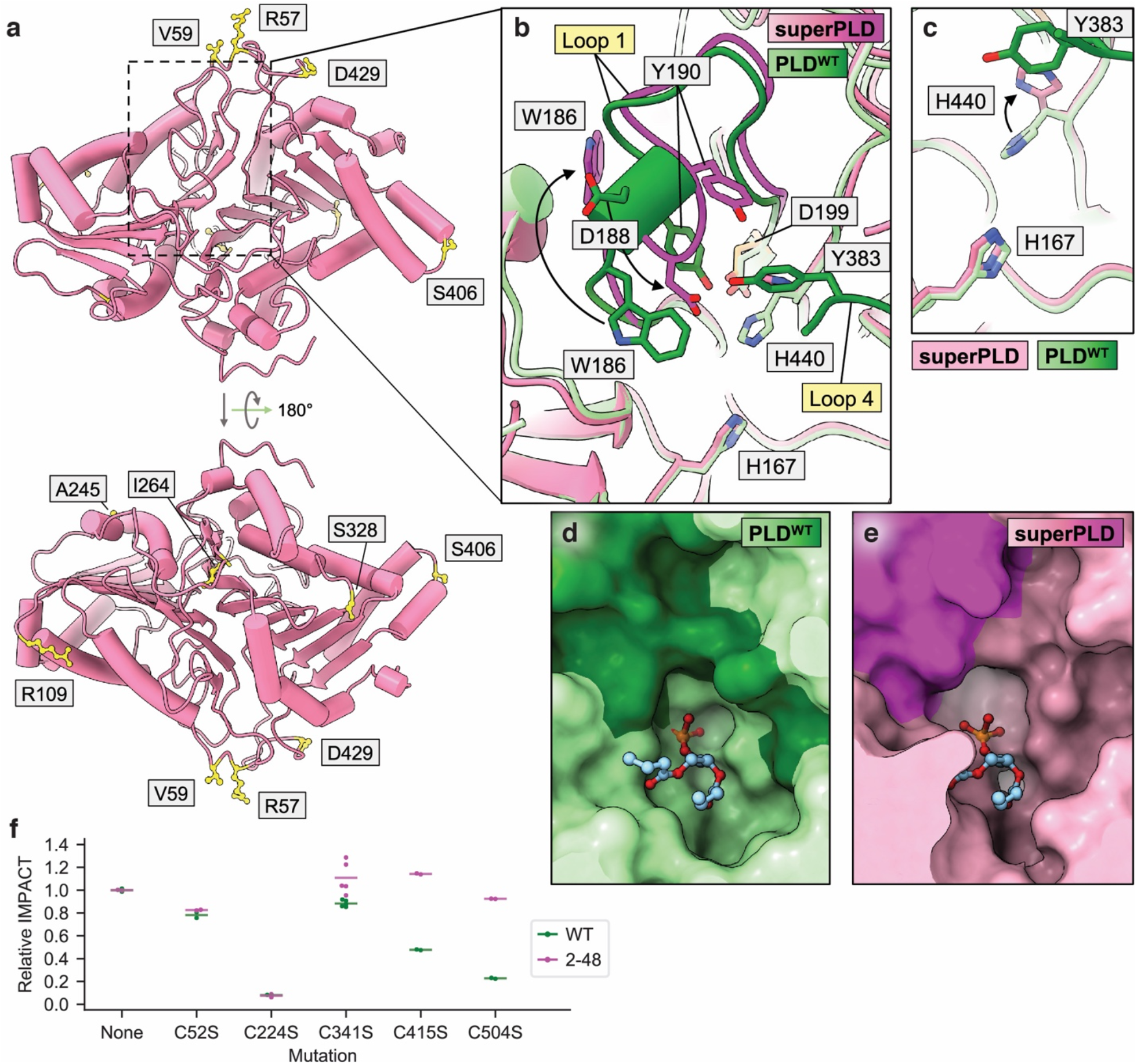
Structural comparison of superPLD and PLD^WT^. **a**, Crystal structures of superPLD (2-48). Residues mutated in superPLD are shown in yellow. **b**, Zoomed-in structures around loops 1 and 4. Structures of PLD^WT^ and superPLD are shown in green and magenta, respectively, with loops 1 and 4 in each structure shown in a darker color. **c**, Zoomed-in structures around His 440. **d–e**, Surface structures around the catalytic pocket of PLD^WT^ (**d**) and superPLD (**e**) highlighting the larger cavity in superPLD. **f**, Comparison of the effect of mutating cysteine residues on PLD^WT^ (green) vs. superPLD (2-48; magenta) to prevent disulfide bond formation. IMPACT fluorescence intensity normalized to optoPLD expression was determined by flow cytometry. Horizontal lines indicate average (n=2–5) of relative mean intensities of IMPACT fluorescence of cells expressing plasma membrane-targeted optogenetic versions of the indicated mutant PLD compared to the parental PLD (e.g., PLD^WT^ or superPLD (2-48)), as measured by flow cytometry.

A second difference between superPLD and PLD^WT^ inside the catalytic pocket was the orientation of H440, which has flipped away from the active site in the superPLD structures relative to its position in the PLD^WT^ structure (Fig. 6c). As H440 directly participates in the reaction mechanism as a general acid/base catalyst (along with the catalytic nucleophile H167)^45,46^, the flipped position of H440 most likely represents the post-catalytic form of superPLD. Consistent with this hypothesis, we found that superPLD co-purified with its product PA, with the electron density of the glycerophosphate head group located in between H167 and H440 but at distances too far from either histidine residue for any productive catalysis (Supplementary Fig. 9a–d). Mutational analysis of superPLD confirmed that residues known to be important for catalysis in PLD^WT^ were equally important in superPLD, indicating that superPLD most likely shares the similar ligand-binding site as PLD^WT^ in its active form (Supplementary Fig. 9e).

Finally, the structural and mutational analysis revealed that superPLD was less reliant upon its four intramolecular disulfide bonds than PLD^WT^. First, the C295–C341 disulfide bond present in PLD^WT^ was absent in superPLD (Supplementary Fig. 10a–d). Second, mutagenesis revealed that superPLD retained its activity much better than PLD^WT^ when the C415–C504 disulfide bond was removed (Fig. 6f). Interestingly, we noticed that the most active superPLDs (2-23 and 2-48) contained two nearby Gly-to-Ser mutations in short unstructured turns or loops adjacent to helices or sheets (G328S and G406S). We postulated that these mutations might favor folding via interstrand contacts, even in the absence of covalent crosslinking provided by the disulfide, due to reduced entropy from a more restricted conformational space (Supplementary Fig. 10e). Indeed, reversion of those residues to Gly within the superPLD background rendered the enzyme equally sensitive to C415S mutation as PLD^WT^ (Supplementary Fig. 10f–g). Collectively, these analyses point to possible sources of the increased cellular activity of superPLDs and highlight the utility of the mammalian cell-based activity-dependent directed evolution platform for generating superPLDs optimized for use for membrane editing in the reductive environment of cytosol-facing membranes in mammalian cells.

## DISCUSSION

We developed an activity-based directed evolution platform in mammalian cells for improving the efficiency and utility of PLD for membrane editing. We achieved the generation of a series of superPLDs with a wide range of activities, with the most active superPLD exhibiting a two orders-of-magnitude activity increase compared to PLD^WT^. SuperPLDs efficiently catalyzed both hydrolysis and transphosphatidylation of PC in cells and *in vitro*, enabling the synthesis of a variety of natural and unnatural phospholipids with designer head groups. SuperPLDs are poised for several types of applications. In cell biology, superPLDs can act as a general membrane editor of the phospholipidome, where products can be dialed in simply by the choice of alcohol substrate^36^. We also envision that recombinant superPLDs will be highly desired to replace unoptimized PLDs isolated from natural sources as industrial biocatalysts for the efficient chemoenzymatic synthesis of phospholipids.

The structural and biochemical studies to examine the sources of improved activity of superPLDs compared to PLD^WT^ underscore the importance of the activity-based directed evolution strategy in mammalian cells that was used to generate the superPLDs. Each superPLD bears 6–10 mutations, which cooperatively increase enzymatic activity. Instead of targeting residues that directly contact the substrates, the mutations caused a reorganization of loops that demarcate the active site opening, leading bulky residues to flip away and create a larger cavity, speaking to the importance of random mutagenesis in our directed evolution strategy. The structural changes in superPLDs also rendered these enzymes more tolerant to disulfide reduction, despite no changes in thermal stability, a likely result of the evolution having been performed in the mammalian cytoplasmic compartment. This latter property also greatly facilitated the purification of superPLDs from *E. coli* in high yield, supporting their potential development into industrial biocatalysts.

Beyond the utility of superPLDs, this study is significant for expanding the scope of directed enzyme evolution in mammalian cells to include activity-based fluorescent labeling for enzyme engineering. Metabolic and chemoenzymatic labeling are popular strategies to probe or perturb various cellular events, including lipid metabolism^7,47^, glycosylation^48–51^, and other post-translational modifications^52–54^. We envision that extension of our mammalian cell-based directed evolution platform to other enzyme classes will enable engineering of highly efficient enzymes capable of serving as editors of a wider set of biomolecules in mammalian cells.

## METHODS

### Mammalian cell culture, transfection, and lentiviral transduction

Cells were grown in DMEM (Corning) supplemented with 10% FBS (Corning), 1% penicillin/streptomycin (Corning), and 1 mM sodium pyruvate (Thermo Fisher) at 37 °C in a 5% CO2 atmosphere. For poly-L-lysine pre-treatment, cell plates were treated with 0.1 mg/mL poly-L-lysine (Sigma Aldrich; P2636) in PBS for 1–12 h at 37 °C, followed by triple rinses with autoclaved deionized water.

For transient transfection, HEK 293T cells were transfected using Lipofectamine 2000 (Invitrogen; 11668019) following the manufacturer’s protocol. Briefly, cells were incubated in regular DMEM media containing plasmids pre-mixed with Lipofectamine 2000 (0.3 μg optoPLD plasmid and 0.75 μL Lipofectamine 2000 per well for a 24-well plate), and the cells were incubated for 20–24 h before being labeled and analyzed.

For lentivirus production, HEK 293TN cells seeded on a 6-well plate were incubated in Transfectagro (Corning) supplemented with 10% FBS containing plasmids pre-mixed with Lipofectamine 2000 (0.5 μg envelope plasmid, 1 μg packaging plasmid, 1.5 μg optoPLD plasmid, and 6 μL Lipofectamine 2000 per well for a 6-well plate). 8 h after transfection, the transfection media was replaced with regular DMEM media, and media were collected 24 h and 48 h after transfection to obtain virus-containing media. For lentiviral transduction, HEK 293T cells seeded on a 6-well plate (pre-treated with poly-L-lysine) were incubated in 1.5 mL virus-containing media supplemented with 0.5 mL fresh media and 0.8 μg/mL polybrene (Millipore Sigma). The 6-well plate was covered with aluminum foil to keep cells in the dark. After 24 h, virus-containing media was replaced with fresh DMEM media, and cells were incubated in the dark for another 24 h before being labeled and sorted (details in “IMPACT labeling and cell sorting” section).

### Generation of optoPLD libraries and mutants

Libraries of optoPLD mutants were generated by error-prone PCR as described previously^55^. Briefly, 100 ng of the template DNA was amplified with 0.5 μM forward and reverse primers (Supplementary Table 2, primers 10 and 11), 200 μM dNTPs mix, 2 μM 8-oxo-dGTP (TriLink BioTechnologies, N-2034), 2 μM dPTP (TriLink BioTechnologies; N-2037), and 2.5 U Taq polymerase in Thermopol Reaction Buffer (New England Biolabs; B9004S). The PCR products were then gel purified and re-amplified for another 25 cycles under normal PCR conditions using the same primers. The second PCR products were digested using BamHI/EcoRI and cloned into optoPLD lentiviral vector (pCDH-CRY2-mCh-PLD-P2A-CIBN-CAAX, Supplementary Table 1, entry 4) digested using the same restriction enzymes. The ligated product was transformed into DH5α *E.coli*, and the grown colonies were scraped and subjected to plasmid extraction. The resulting optoPLD plasmids were used to transfect HEK 293TN cells for lentivirus production (details in “Mammalian cell culture, transfection, and lentiviral transduction” section).

For introducing site-specific mutations to PLD, N-terminal and C-terminal fragments of PLD were amplified using primer 10 (Supplementary Table 2) and a reverse mutagenizing primer containing a desired mutation (Supplementary Table 3) for the N-terminal fragment, and primer 11 (Supplementary Table 2) and a forward mutagenizing primer (Supplementary Table 3) for the C-terminal fragment. The two fragments were then stitched together using overlap-extension PCR to obtain the mutagenized PLD, which was subsequently cloned into an optoPLD transient expression vector (pCDNA3-CRY2-mCh-PLD-P2A-CIBN-CAAX, Supplementary Table 1, entry 6) using BamHI and EcoRI cut sites.

### Setup for optogenetics experiments

A homemade light box was built by attaching four strips of dimmable, 12 V blue-LED tape light (1000Bulbs.com; 2835–60-IP65-B1203) on the inside of a Styrofoam box. For optogenetics experiments, the light box was placed inside the CO2 incubator using an AC Outlet Power Bank (Omars; 24,000 mAh, 80 W) as a power supply. An outlet timer (BN-LINK) was used to switch the light on and off automatically to enable 3-s intervals of blue light in every 1 min.

### IMPACT labeling and cell sorting

PLD1/2 double knockout HEK 293T cells^18^ expressing optoPLD libraries were treated with 1–5 mM azidopropanol for 30 min at 37 °C in the presence of intermittent blue light illumination (3-s pulses every 1 min). After three rinses with PBS, cells were treated with 1 μM bicyclononyne-BODIPY fluorophore (BCN-BODIPY^56^) for 10 min at 37 °C. Cells were again rinsed three times with PBS and incubated in DMEM media for 10 min at 37 °C to remove excess fluorophore. Cells were then trypsinized, resuspended in PBS, and sorted using a Sony MA900 Cell Sorter or a FACSAria Fusion Cell Sorter. Cells expressing optoPLD^dead^, a catalytically dead mutant (H167A), were similarly labeled and sorted as a negative control, and the population in cells expressing optoPLD libraries that showed higher signal than the negative control was collected. The collected cells were expanded, at which point cells were reseeded for another round of selection or subjected to genomic extraction. Flow cytometry plots and histograms showing sorting strategy and collected cell populations for each round of selection are shown in Supplementary Fig. 2.

### Genomic extraction and amplification of PLD fragments

Genomic DNA was extracted from HEK 293T cells using a NucleoSpin Blood kit (Takara Bio; 740951) following the manufacturer’s protocol. Briefly, cells were rinsed once with PBS and lysed, then the lysis was applied to the DNA-binding column. After rinsing and drying the column, 60 μL of water was applied to elute DNA. The eluate was used as a template for PCR reactions to amplify PLD fragments. For PCR reactions, 0.5–10 μL of template was amplified for 25 cycles under normal PCR conditions with primers 8 and 9 (Supplementary Table 2, for use with Taq polymerase) or primers 10 and 11 (Supplementary Table 2, for use with Phusion polymerase). The PCR products were digested and cloned into the optoPLD vector as described in “Generation of optoPLD libraries” section.

### Directed evolution of optoPLDs

For the first round of evolution, two optoPLD libraries were generated using PLD^WT^ or PLD^G429D^ (the G429D mutation exhibits modestly higher (~1.3-fold) activity than PLD^WT^)^20^ as the starting template. The optoPLD libraries were introduced into HEK 293T cells using lentiviral transduction, and cells expressing optoPLD libraries were labeled and sorted as described above in “IMPACT labeling and cell sorting” section. The sorted cells were expanded prior to another cycle of IMPACT labeling and cell sorting. After the second cycle of selection, cells were subjected to genomic extraction. PLD fragments were amplified from the extracted DNA using Taq polymerase to introduce more mutations and then cloned into optoPLD vector.

For the second and third rounds of evolution, the optoPLD libraries were generated using the product of the previous round of evolution as the template. For these rounds, Taq polymerase, which has lower fidelity and thus expected to introduce ~1 mutation per PLD, was used for amplification. Further mutations were added by error-prone PCR, and the libraries with and without error-prone PCR were combined. The generated optoPLD libraries were expressed in cells, and cells were labeled and sorted as described above. The sorted cells were subjected to two more cycles of selection, followed by genomic extraction and PLD amplification.

For the fourth and subsequent rounds of evolution, genomic DNA extracted from cells was amplified by Phusion polymerase to minimize the introduction of further mutations. The rest of the evolution was performed likewise.

After evolution, PLD mutants were cloned into an optoPLD transient expression vector (Supplementary Table 1, entry 6) using BamHI and EcoRI cut sites. Each plasmid isolated from a single *E.coli* colony was analyzed by Sanger sequencing using primers 28 and 29 (Supplementary Table 2) to determine the mutations in each clone of PLD mutants.

### Quantitative comparison of PLD activity using IMPACT

HEK 293T cells were transiently transfected with CRY2-mCh-PLD-P2A-CIBN-CAAX, where the PLD sequence contained indicated set of mutations, and cells were kept in dark for 18–24 h. For IMPACT labeling, cells were treated with 0.1–1 mM azidopropanol for 30 min at 37 °C in the presence of intermittent blue light illumination (3-s pulses every 1 min). After three rinses with PBS, cells were treated with 1 μM BCN-BODIPY for 10 min at 37 °C, again rinsed three times with PBS, and incubated in DMEM media for 10 min at 37 °C. Cells were then trypsinized and subjected to flow cytometry analysis using an Attune NxT flow cytometer to measure mCherry and BODIPY fluorescence, which correspond to optoPLD expression level and IMPACT labeling intensity, respectively. Cells expressing similar amounts of optoPLD were gated, and the average IMPACT signal in the gated population was used to compare PLD activity of different mutants (Supplementary Fig. 2t).

### Evaluation of phosphatidic acid localization by confocal microscopy

Due to the large DNA size of CRY2-mCherry-PLD-P2A-CIBN-CAAX, which affected lentivirus production efficiency, CRY2-mCherry-PLD and CIBN-CAAX were packaged separately into lentivirus. pCDH-CRY2-mCherry-superPLD was prepared by cloning superPLD into an optoPLD lentiviral expression vector (Supplementary Table 1, entry 8) using BamHI and EcoRI cut sites. Lentivirus containing GFP-PASS, CRY2-mCherry-PLD, and CIBN-CAAX (for plasma membrane-targeted optoPLD) or p18-CIBN (lysosome-targeted optoPLD) were prepared as described in “Mammalian cell culture, transfection, and lentiviral transduction” section. Spinfection was used for efficient co-transduction of HEK 293T cells with the three lentivirus preparations. Briefly, cells were seeded on 35-mm glass-bottom imaging dishes (Matsunami Glass), and after the addition of lentivirus-containing media to cells, cells were centrifuged at 931 g for 2 h at 37 °C. After spinfection, lentivirus-containing media was replaced with regular growth media, and cells were kept in the dark for 48 h before imaging.

For colocalization analysis with LysoView 633, HEK 293T cells transduced with GFP-PASS, CRY2-mCherry-PLD and p18-CIBN were prepared as described above, and 1X LysoView 633 was added before the imaging. For evaluation of LKB1 localization, HEK 293T cells seeded on imaging dishes were transfected with GFP-LKB1 and p18-CIBN-P2A-CRY2-mCherry-PLD using Lipofectamine 2000, and cells were kept in the dark for 20 h before imaging.

Images were acquired every 1 min for 1 h at 37 °C using Zeiss Zen Blue 2.3 on a Zeiss LSM 800 confocal laser scanning microscope equipped with Plan Apochromat objectives (40X 1.4 NA) and two GaAsP PMT detectors. Solid-state lasers (488, 561, and 640 nm) were used to excite GFP, mCherry, and LysoView 633, respectively, and the 488 nm laser irradiation also served as a stimulus for activating optoPLD recruitment to the plasma membrane or lysosomes. The colocalization between GFP-PASS/LKB1 and CRY2-mCherry-PLD was calculated for each transfected cell using Coloc 2 plugin on ImageJ.

### PLD purification using affinity and size-exclusion chromatography

PLD construct was cloned into the pCAV4.1 vector^57^ (Supplementary Table 1, entry 13), which is a modified T7 expression vector containing an N-terminal 6xHis-NusA tag followed by peptide sequence that is cleavable by the HRV 3C protease. Constructs were transformed into *Rosetta 2* (DE3) or *Rosetta-gami 2* (DE3) *E. coli*, grown at 37 °C in 2 x 1 L terrific broth media supplemented with chloramphenicol (25 μg/mL) and ampicillin (100 μg/mL) to an OD600 of 0.8, and then induced with IPTG (0.1 mM) for 20 h at 18 °C. Cells were harvested by centrifugation, resuspended in 50 mL bacterial lysis buffer (50 mM sodium phosphate, 500 mM NaCl, 10% glycerol, pH 7.5) supplemented with 5 mM β-mercaptoethanol (for *Rosetta)* and 0.5 mM phenylmethylsulfonyl fluoride (PMSF), and homogenized by using a Sonic Dismembrator (Fisherbrand Model 505). The cell lysate was centrifuged at 30,000 g for 30 min, and the supernatant was incubated with 2 mL TALON Metal Affinity Resin (Takara Bio) for 1 h at 4 °C with rotation. The resin was then loaded onto a disposable column (Bio-Rad) and rinsed for 5 times with 25 mL bacterial lysis buffer. After resuspending the washed resin in 5 mL bacterial lysis buffer, His-tagged HRV3C protease was added and the mixture was incubated overnight at 4 °C with rotation to elute PLD from the resin. The supernatant containing cleaved PLD was concentrated using an Amicon 0.5 mL 10 kDa molecular weight cutoff centrifugal filter. For crystallization and thermal stability analysis, further purification of PLD using size-exclusion chromatography was performed using an ÄKTA pure system equipped with a Superdex 200 Increase 10/300 GL column in 20 mM Tris-HCl (pH 8.0) and 150 mM NaCl.

### *In vitro* kinetics assays of PLD activity

PLD activity was determined using the Amplex Red Phospholipase D Assay Kit following the manufacturer’s protocol. Briefly, 100 μM Amplex Red, 2 U/mL horseradish peroxidase (HRP), 0.2 U/mL choline oxidase, and 0.02–0.4 mg/mL 1,2-dioleoyl-*sn*-glycero-3-phosphatidylcholine (DOPC; prepared 40 mg/mL in ethanol) were added to PBS (pH 7.4) to prepare a master mix solution. The solution was added to 10 ng/mL PLD to start the reaction, and fluorescence signal was measured during the incubation at 37 °C using a BioTek Synergy H1 Microplate Reader. The luminescence signal at the reaction endpoint, when all the DOPC was consumed, was used to convert luminescence signal (AU) to [product] (μM) for calculating *V*_max_. *K*_m_ and *k*_max_ of the reaction were calculated based on the Michaelis-Menten equation.

### Phospholipid synthesis and LC–MS analysis

In 1.5-mL Safe-Lock Eppendorf tubes, 50 μM–2 M of alcohol and 0.1 μg of PLD were added to 100 μL PBS (pH 7.4). For ethanolamine, the pH was adjusted to pH 7.4 by addition of HCl. After addition of 0.8 mg of DOPC in 80 μL ethyl acetate, the tubes were placed in a plastic box and shaken vigorously in a 37 °C shaker for 1–24 h at 350 rpm. The reaction was quenched by adding 250 μL methanol, 125 μL acetic acid (20 mM in water), and 500 μL chloroform. The solution was mixed thoroughly by shaking vigorously for 5 min, and the tubes were centrifuged at high speed for 1 min. 10 μL aliquots of the bottom organic layer were collected and transferred into new tubes. Solutions were diluted, filtered, and subjected to high-resolution LC–MS analysis to quantify the concentrations of DOPC, dioleoyl phosphatidic acid (DOPA), and dioleoyl phosphatidyl alcohol (DOPAlc) in the sample. The obtained concentration was used to calculate the total amount of each compound in the reaction mixture, which was used to determine the percent yield for DOPA and DOPAlc.

LC–MS analysis was performed on an Agilent 6230 electrospray ionization–time-of-flight MS coupled to an Agilent 1260 HPLC equipped with a Luna 3 μm Silica LC Column (Phenomenex; 50 × 2 mm) using a binary gradient elution system where solvent A was chloroform/methanol/ammonium hydroxide (85:15:0.5) and solvent B was chloroform/methanol/water/ammonium hydroxide (60:34:5:0.5). Separation was achieved using a linear gradient from 100% A to 100% B over 10 min. Phospholipid species were detected using an Agilent Jet Stream source operating in positive or negative mode, acquiring in extended dynamic range from m/z 100–1700 at one spectrum per second; gas temperature: 325 °C; drying gas 12 L/min; nebulizer: 35 psig; fragmentor 300 V (for positive mode) and 250 V (for negative mode); sheath gas flow 12 L/min; Vcap 3000 V; nozzle voltage 500 V.

### Thermal stability analysis

The thermal stability of PLD^WT^ and superPLDs was determined as previously reported^58^. Briefly, PLD^WT^ and superPLDs were diluted to 0.1 mg/mL final concentration in Tris-HCl buffer (10 mM, pH 8.0, 150 mM NaCl) containing SYPRO Orange (1:1000 dilution of 5000X concentrate). The fluorescence signal was measured while the temperature was slowly raised using a Roche LightCycler 480. Melting temperature (Tm) was determined by the temperature at which the fluorescence signal reached at 50% of its maximum.

### Chemical stability analysis

The chemical stabilities of PLD^WT^ and superPLD (2-48) were determined by measuring the residual activity of PLD treated with urea. 1 μg/mL PLD was incubated in solutions of 0–4 M urea in PBS for 12 h at 37 °C, after which the PLD activity was measured as described in “In vitro kinetics assays of PLD activity” section. The relative rate of reaction compared to 0 M urea (untreated) PLD was used for estimating the chemical stability.

### Evaluation of YAP localization by immunofluorescence

HEK 293T cells seeded on cover glasses coated with poly-L-lysine were transfected with CRY2-mCherry-PLD-P2A-CIBN-CAAX, and cells were kept in dark for 16 h before being placed in a serum-starvation medium (DMEM supplemented with 1% penicillin/streptomycin without FBS). After 6 h of starvation, cells were stimulated for 1 h with intermittent blue light illumination (5-s pulses every 1 min), followed by cell fixation and immunostaining as described previously^20^. Briefly, cells were fixed in 4% paraformaldehyde for 10 min at room temperature, followed by extraction in a solution of 0.5% Triton X-100 in PBS for 5 min. Cells were then blocked in a solution of 1% BSA and 0.1% Tween-20 in PBS (blocking buffer) for 30 min. Immunostaining was then performed by treating cells with a 1:100 dilution of anti-YAP antibody (Santa Cruz Biotechnology; sc-101199) in blocking buffer for 1 h, rinsing three times with 0.1% Tween-20 in PBS solution (PBS-T), treatment with a 1:1,000 dilution of anti-mouse–Alexa Fluor 488 antibody conjugate (Invitrogen; A-21202) in blocking buffer for 1 h, and rinsing three times with PBS-T. Cells were mounted on microscope slides using ProLong Diamond Antifade Mountant with DAPI (Thermo Fisher) and incubated overnight at room temperature in the dark. Image acquisition by laser-scanning confocal microscopy was performed as described above by using solid-state lasers (405, 488, and 561 nm) to excite DAPI, Alexa Fluor 488, and mCherry, respectively.

### Quantification of p-AMPK and p-S6K by Western blotting

HEK 293T cells were transduced with CRY2-mCherry-PLD and CIBN-CAAX using lentivirus and spinfection as described in “Evaluation of phosphatidic acid localization by confocal microscopy” section. Cells were incubated with either 10 μM STO-609 (CaMKK inhibitor; for AMPK signaling assay) for 6 h or 10 μM dorsomorphin (AMPK inhibitor; for mTOR signaling assay) for 1 h at 37 °C, followed by 30 min stimulation with intermittent blue light illumination (5-s pulses every 1 min). Cells were then lysed with RIPA lysis buffer supplemented with protease and phosphatase inhibitors (50 mM Tris-HCl, pH 7.4, 150 mM NaCl, 1% Triton-X, 0.5% sodium deoxycholate, 0.1% SDS, 1 mM EDTA, 1x cOmplete™ Protease Inhibitor, 17.5 mM betaglycerophosphate, 20 mM sodium fluoride, 1 mM activated sodium orthovanadate, 5 mM sodium pyrophosphate). After sonication and centrifugation, the lysate supernatants were mixed with 6x Laemmli sample buffer to prepare the sample for Western blotting. The membrane was blotted with antibodies for phospho-AMPKα (Thr172) (Cell Signaling Technology; #2535), phospho-p70 S6 kinase (Thr389) (Cell Signaling Technology; #9205), p70 S6 kinase (Santa Cruz Biotechnology; sc-8418), mCherry (Novus Biologicals; NBP1-96752), or actin (MP Biomedicals; 08691001), with detection by chemiluminescence using the Clarity Western ECL Substrate (Bio-Rad) and acquisition on a Bio-Rad ChemiDoc MP System.

### Quantification of substrate conversion by superPLD in cells

HEK 293T cells seeded on 12-well plates were transduced with CRY2-mCherry-PLD and CIBN-CAAX using lentivirus and spinfection as described above. Cells were incubated with 0.5– 2% ethanol, which should be sufficient to inhibit most of PLD hydrolysis activity^22^, for 30 min with intermittent blue light illumination. Cells were then rinsed with PBS three times and subjected to lipid extraction. For lipid extraction, cells were scraped in 250 μL methanol, 125 μL acetic acid (20 mM in water), and 100 μL PBS. The cell suspension was transferred into a 1.5-mL centrifuge tube, and the lipids were extracted and subjected to LC–MS analysis as described in “Phospholipid synthesis and LC–MS analysis” section.

### Crystallization of superPLD

Crystals for superPLD (2-48 mutant) were obtained by mixing 1 μL of purified protein at 2.5 mg/mL with 1 μL of well solution containing 21% PEG, 0.15 M Li_2_SO_4_, and citrate-NaOH (pH 4.4) and equilibrated against 200 μL of well solution at 18 °C. Crystals grew within 5–7 d. Single crystals were harvested and soaked in the well solution supplemented with 10% ethylene glycerol for 10 s before plunge freezing in liquid N2. Crystals for the 2-23 mutant were obtained by mixing 1 μL of purified protein at 2.5 mg/mL with 1 μL of well solution containing 19% PEG, 0.15 M Li2SO4, citrate-NaOH (pH 4.15) and equilibrated against 200 μL of well solution at 18 °C. Crystals grew within 5–7 d. Single crystals were harvested and soaked in the well solution supplemented with 10% ethylene glycol for 10 s before plunge freezing in liquid N2.

### X-ray diffraction data collection, processing and model building

Diffraction experiments were conducted at beamline 24-ID-E of the Advanced Photon Source (APS) and beamline ID7B2 of the Cornell High Energy Synchrotron Source (CHESS). Diffraction data sets were collected at 100 K and processed using XDS^59^. Crystals of superPLD (2-48 mutant) and 2-23 mutant diffracted to 1.85 Å and 1.9 Å, respectively. The crystal structure of PLD^WT^ (PDB ID 1V0Y) was used to obtain phasing information using molecular replacement using Phaser^60^ in PHENIX^61^. The models were subjected to iterative rounds of manual re-building using COOT^62^ followed by refinement in PHENIX^61^. We observed electron density in the active site that likely corresponds to a bound reaction intermediate. Based on previously reported structures of PLD in complex with reaction intermediates (PDB 7JRU and 7JRV), we modelled a phosphate moiety in part of this density. We note that the density was not clear enough to model the glycerol back bone and the acyl-chains, so we omitted these groups from the model. Final refinement and validation statistics for the models are reported in Supplementary Table 4. Structural models and structure factors will be available in the RCSB PDB upon publication.

### Statistical analysis

Statistical significance was calculated using one-way ANOVA, followed by Tukey’s HSD test using the “statsmodel” Python package. *, *p* < 0.05; **, *p* < 0.01; ***, *p* < 0.001.

## ACKNOWLEDGMENTS

J.M.B. acknowledges support from a Beckman Young Investigator award, a Sloan Research Fellowship, and the NSF (CAREER CHE-1749919). R.T. was supported by Honjo International, Funai Overseas, and Cornell Fellowships. J.C.F. and S.R.B were supported by NIH/NIGMS grant R35GM136358. Beamline 24-ID-E of APS is funded by the National Institute of General Medical Sciences from the National Institutes of Health (P30 GM124165). The Eiger 16M detector on the 24-ID-E beam line is funded by a NIH-ORIP HEI grant (S10OD021527). This research used resources of the Advanced Photon Source, a U.S. Department of Energy (DOE) Office of Science User Facility operated for the DOE Office of Science by Argonne National Laboratory under Contract No. DE-AC02-06CH11357. Beamline ID7B2 of CHESS is supported by the National Science Foundation under award DMR-1829070, and by award 1-P30- GM124166-01A1 from the National Institute of General Medical Sciences, National Institutes of Health, and by New York State’s Empire State Development Corporation (NYSTAR).

## AUTHOR CONTRIBUTIONS

R.T. and J.M.B. designed the study and analyzed data; R.T. carried out directed evolution, molecular cloning, in cellular and *in vitro* activity assays, protein production, and purification;

S.R.B. and R.T. performed protein crystallization and X-ray data collection; S.R.B. analyzed structural data; J.C.F. supervised X-ray crystallography analysis; R.T. and J.M.B. wrote the manuscript, with input from all authors.

## Notes

### Competing Interest Statement

The authors have declared no competing interest.

