## Supplementary Information for "Activity-based directed evolution of a membrane editor in mammalian cells"

### Supplementary Figures

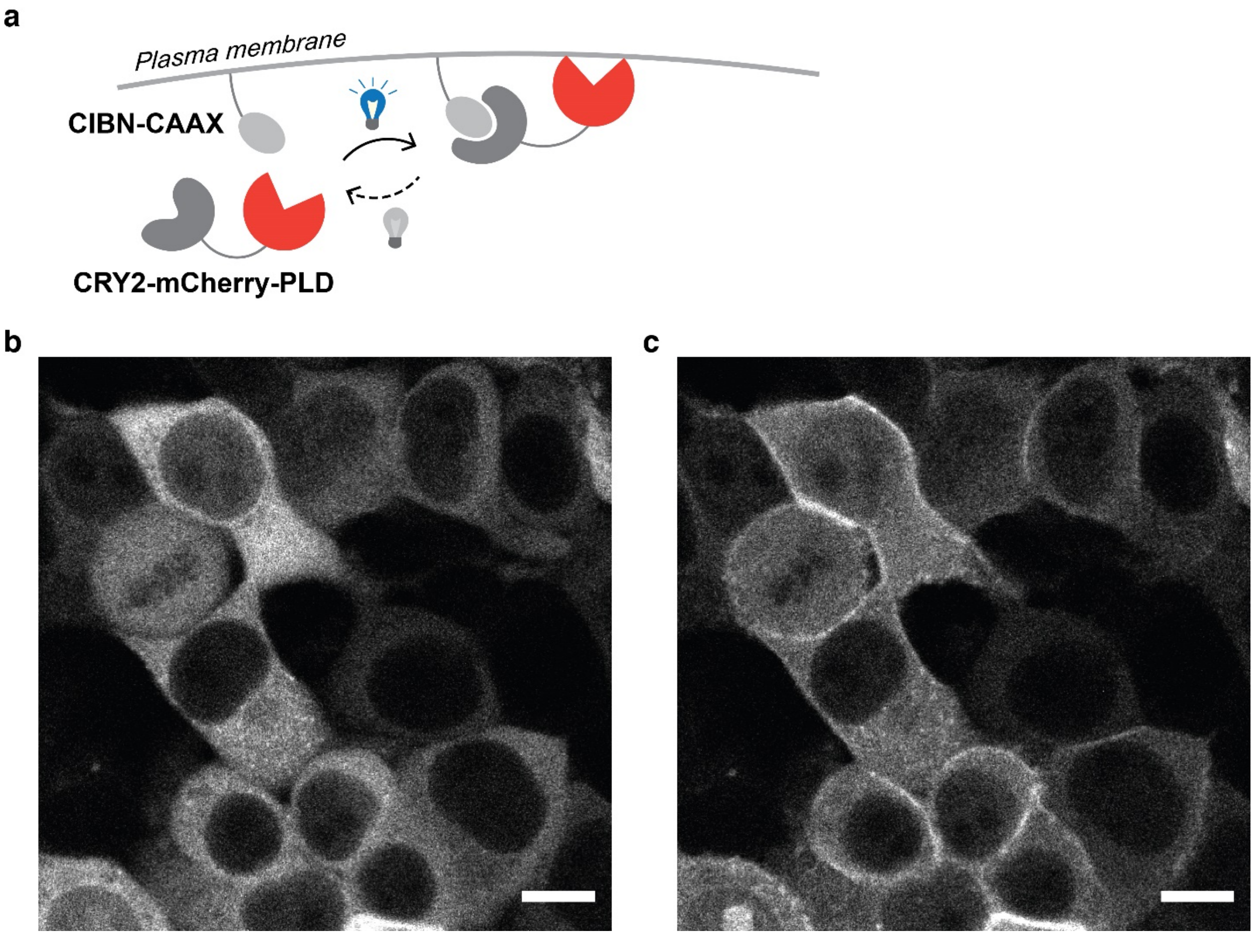

**Supplementary Figure 1. A plasma membrane-targeted optoPLD library. a**, Schematic depicting the design of plasma membrane-targeted optoPLD, which consists of CRY2-mCherry-PLD and CIBN-CAAX. OptoPLD-targeting membrane can be controlled by swapping the plasma membrane-targeted domain, CAAX, with another membrane-targeting domain. **b–c**, Confocal microscopy images of mCherry fluorescence showing localization of optoPLD library (after FACS-based selection) in HEK 293T cells before (**b**) and after (**c**) illumination with 488 nm light. Scale bar: 10 µm.

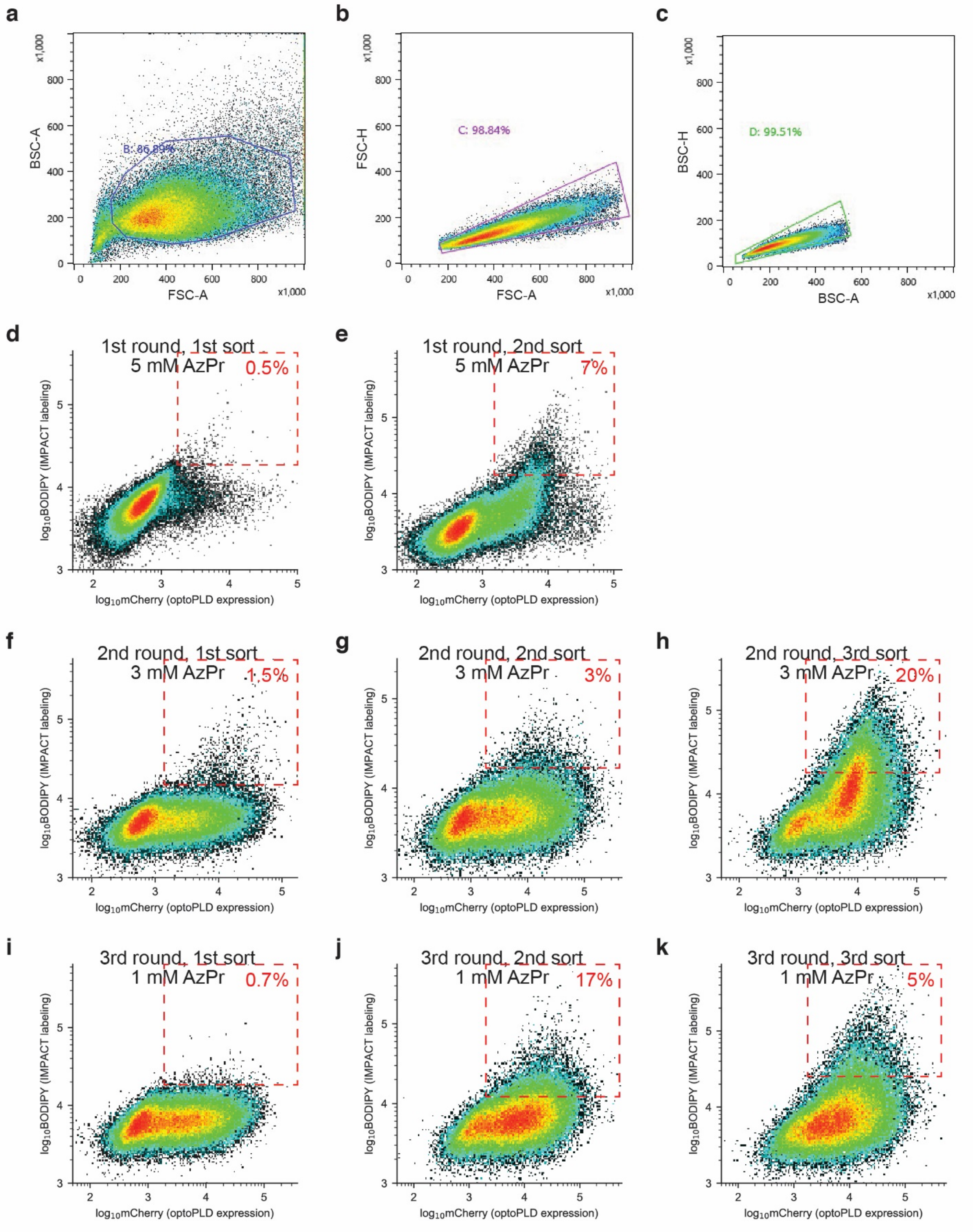
a
b
c
BSC-A
FSC-A
FSC-H
BSC-H
x1,000
B: 86.89%
C: 98.84%
D: 99.51%
d
1st round, 1st sort
5 mM AzPr
0.5%
e
1st round, 2nd sort
5 mM AzPr
7%
f
2nd round, 1st sort
3 mM AzPr
1.5%
g
2nd round, 2nd sort
3 mM AzPr
3%
h
2nd round, 3rd sort
3 mM AzPr
20%
i
3rd round, 1st sort
1 mM AzPr
0.7%
j
3rd round, 2nd sort
1 mM AzPr
17%
k
3rd round, 3rd sort
1 mM AzPr
5%
log10BODIPY (IMPACT labeling)
log10mCherry (optoPLD expression)

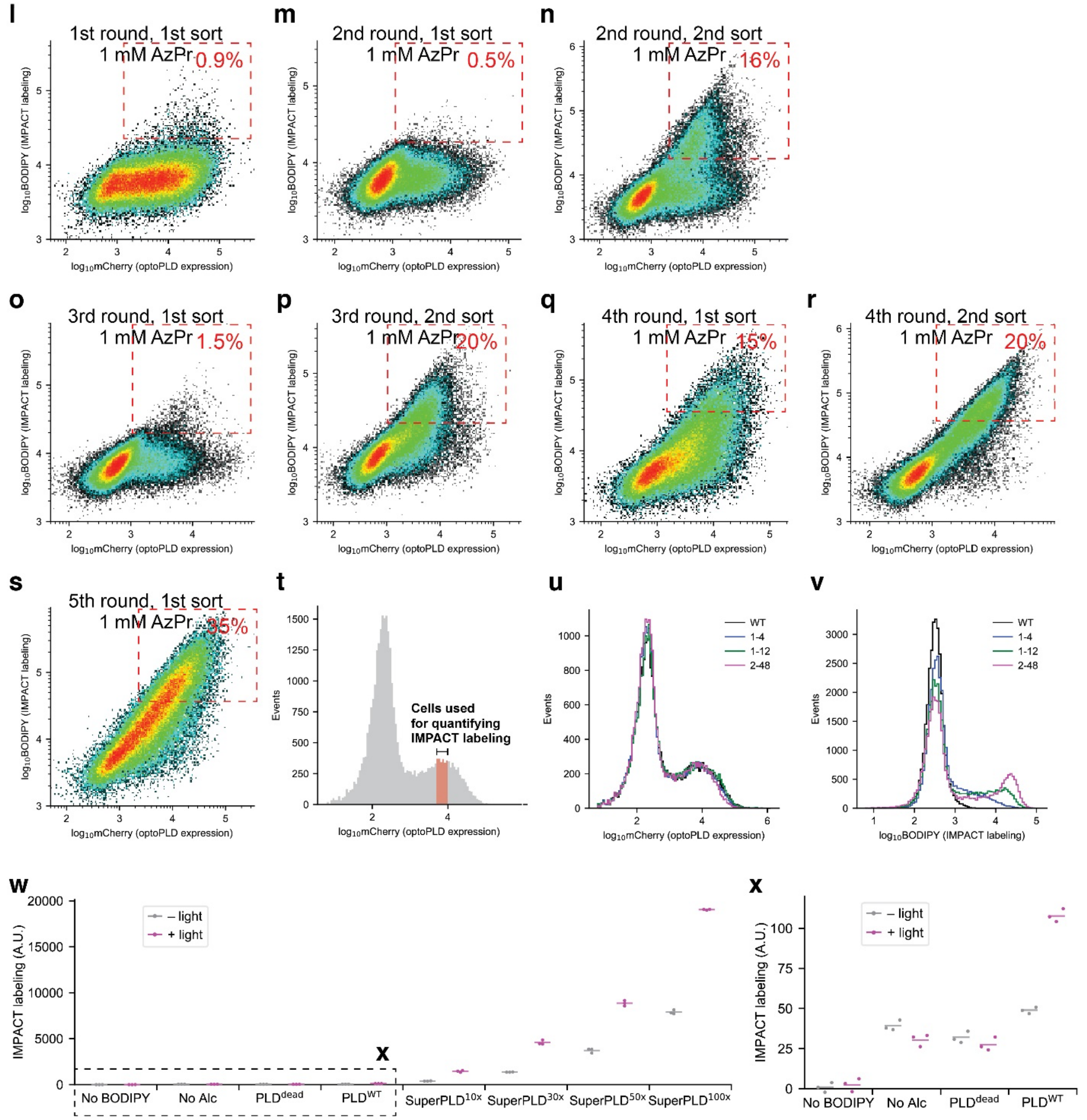

**Supplementary Figure 2. FACS plot and gating strategy for each sorting of directed evolution and superPLD characterization by IMPACT. a–c**, Gating strategy for sorting and analysis of HEK 293T cells expressing optoPLD mutants labeled by IMPACT. Plots of FSC-A vs. BSC-A, population B selected (**a**), FSC-A vs. FSC-H, population C selected (**b**), and BSC-A vs. BSC-H, population D selected (**c**) were used to gate for live, singlet cells. **d–k**, FACS plot of PLD library from each sorting in three rounds of selection with mutagenesis. **l–s**, FACS plot of PLD library from each sorting in five rounds of selection without mutagenesis. The round number, sort number within that round, concentration of azidopropanol (AzPr) used for IMPACT labeling, and the percentage of cells collected are indicated in each plot. **t**, Gating strategy for quantitative comparison of IMPACT labeling. The average BODIPY signal of cells with similar amount of mCherry signal (population shown in red, which is gated for an mCherry fluorescence of $5x10^3$–

$1x10^4$), was used. **u–v**, mCherry (**u**) and IMPACT labeling (**v**) histograms of cells expressing $PLD^{WT}$ (black), 1-4 (blue), 1-12 (green) and 2-48 (magenta), demonstrating that optoPLD mutants with different activity have similar expression levels. **w–x**, The effect of light stimuli on optoPLD activity. Cells expressing optoPLD were treated with 0.5 mM azidopropanol with or without intermittent blue light illumination, followed by treatment with 1 μM BCN-BODIPY, and IMPACT fluorescence intensity normalized to optoPLD expression was determined by flow cytometry. Horizontal lines indicate average (n=3) of mean intensities of IMPACT fluorescence. $PLD^{dead}$, a catalytically dead PLD bearing the H167A mutation; $PLD^{WT}$, wild-type PLD; $superPLD^{x10}$, superPLD mutant clone 1-4; $superPLD^{x30}$, superPLD mutant clone 1-12; $superPLD^{x50}$, superPLD mutant clone 1-27; $superPLD^{x100}$, superPLD mutant clone 2-48.

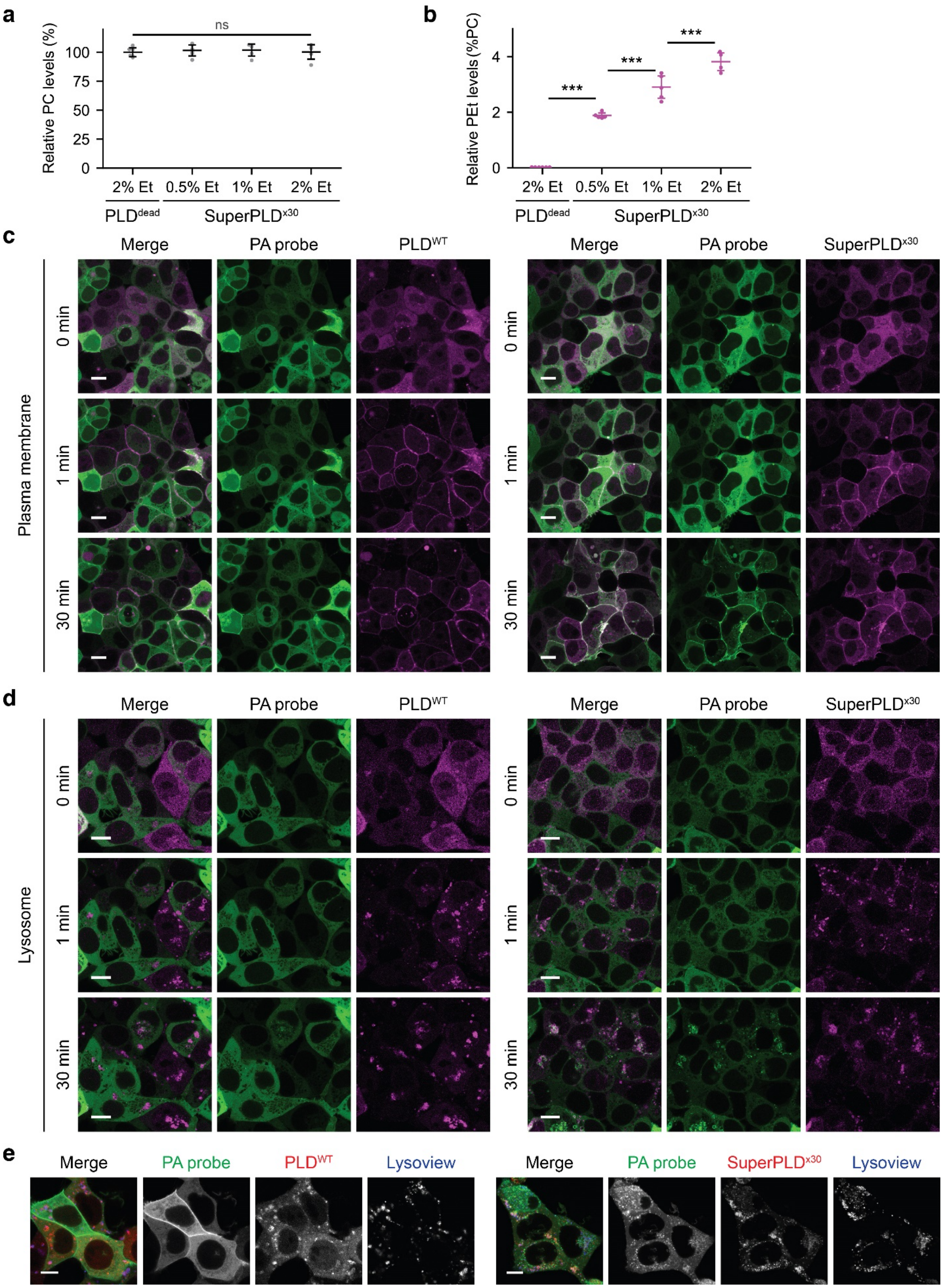
a
Relative PC levels (%)
ns
0
25
50
75
100
2% Et
0.5% Et
1% Et
2% Et
PLDdead
SuperPLDx30
b
Relative PEt levels (%PC)
***
0
2
4
c
Merge
PA probe
PLDWT
SuperPLDx30
Plasma membrane
0 min
1 min
30 min
d
Lysosome
e
Lysoview

**Supplementary Figure 3. Characterization of superPLD activity in cells. a–b**, Quantification of substrate conversion by superPLD in cells. HEK 293T cells expressing plasma membrane-targeted optoPLD (superPLD$^{x30}$) were treated with 0.5–2% ethanol (Et) for 30 min with intermittent blue light illumination. As a control, cells expressing PLD$^{dead}$ were treated with 2% ethanol accordingly. The relative levels of the most abundant PLD substrate (16:0/18:1 PC; POPC) and its transphosphatidylation product (16:0/18:1 PEt; POPEt) were quantified by LC–MS. Relative PC levels compared to PC levels in control samples (PLD$^{dead}$-expressing cells) (**a**) and relative PEt levels compared to PC levels (**b**) are plotted. Horizontal lines indicate average, and vertical error bars indicate standard deviation (n=4–6). **c–d**, Confocal microscopy images of HEK 293T cells co-expressing a PA probe (GFP-PASS) and optoPLD targeted to the plasma membrane (**c**) or lysosomes (**d**) before (0 min), immediately after (1 min), and 30 min after incubation with 488 nm light. **e**, Confocal microscopy images of cells co-expressing the PA probe and the indicated optoPLD construct (WT or superPLD$^{x30}$) targeted to lysosomes, stained with LysoView 633. Scale bar: 10 μm.

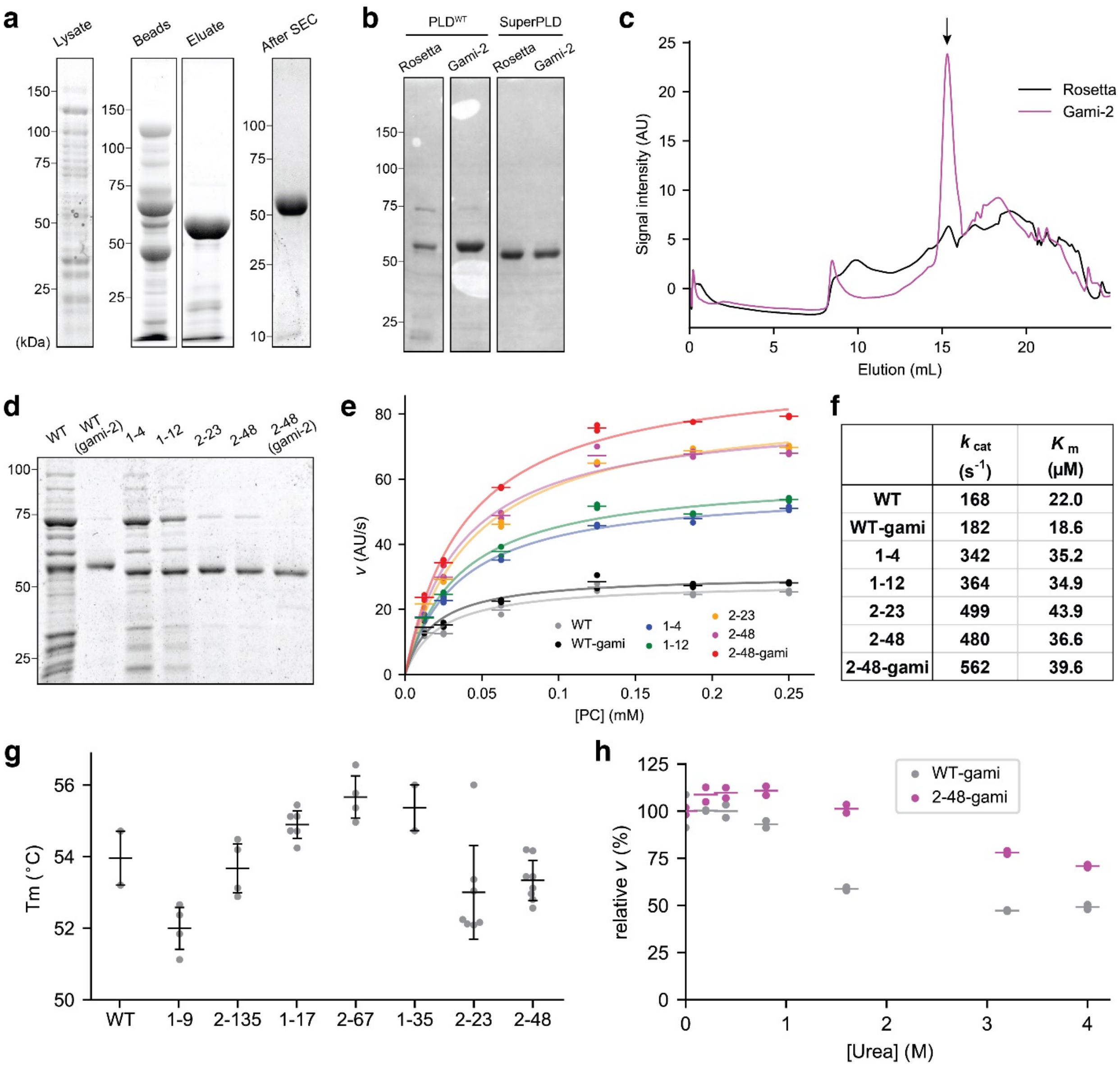

| | $k_{cat}$ ($s^{-1}$) | $K_m$ (μM) |
|---|---|---|
| **WT** | **168** | **22.0** |
| **WT-gami** | **182** | **18.6** |
| **1-4** | **342** | **35.2** |
| **1-12** | **364** | **34.9** |
| **2-23** | **499** | **43.9** |
| **2-48** | **480** | **36.6** |
| **2-48-gami** | **562** | **39.6** |

**Supplementary Figure 4. Purification and in vitro characterization of superPLD and PLD$^{WT}$.** **a**, SDS-PAGE showing His-NusA-superPLD (shown here is clone 2-48) (113 kDa) in cell lysate, His-NusA (58 kDa) being retained on the TALON beads, and superPLD (55 kDa) eluting from the beads and exhibiting high purify after size-exclusion chromatography (SEC). 6xHis-NusA-superPLD was expressed in *E.coli Rosetta 2* and purified using TALON resin. HRV 3C protease was used to cleave between NusA and PLD. **b**, Ponceau stain with PLD$^{WT}$ and superPLD purified from *Rosetta 2* vs. *Gami-2* (*Rosetta-gami 2*) strains. **c**, Size-exclusion chromatogram of PLD$^{WT}$ purified from *Rosetta* vs. *Gami-2*. Black arrow indicates the peak corresponding to PLD. When expressed in the *Rosetta* strain, PLD$^{WT}$ showed significant degradation and lower yield. **d–f**, Kinetic analysis of PLD activity. Equal amounts of PLD were prepared based on SDS-PAGE (**d**) and incubated with indicated concentration of DOPC. Activity assays were performed using the Amplex Red Phospholipase D Assay Kit. The rates of reaction are plotted against the substrate concentration (**e**), and the data were fit to the Michaelis-Menten equation to obtain the kinetic parameters (**f**). **g**, Thermal stability of PLD$^{WT}$ and a subset of superPLD mutants spanning a wide range of activities (see Fig. 2d), indicating no clear correlation between thermal stability and PLD activity. Melting temperature (Tm) was determined in a real-time thermocycler using SYPRO Orange dye. Horizontal lines represent the mean and error bars represent the standard deviation (n=3–8 distinct samples). **h**, Chemical stability of $PLD^{WT}$ and superPLD (2-48) purified from the *Gami-2* strain, indicating increased chemical stability for superPLD. Each enzyme was incubated with indicated concentration of urea in PBS for 12 h at 37 ºC, followed by an activity assay using the Amplex Red Phospholipase D Assay Kit. Relative rates of reaction compared to control samples (enzymes incubated in PBS only) are plotted, and horizontal lines represent the mean (n=2 distinct samples).

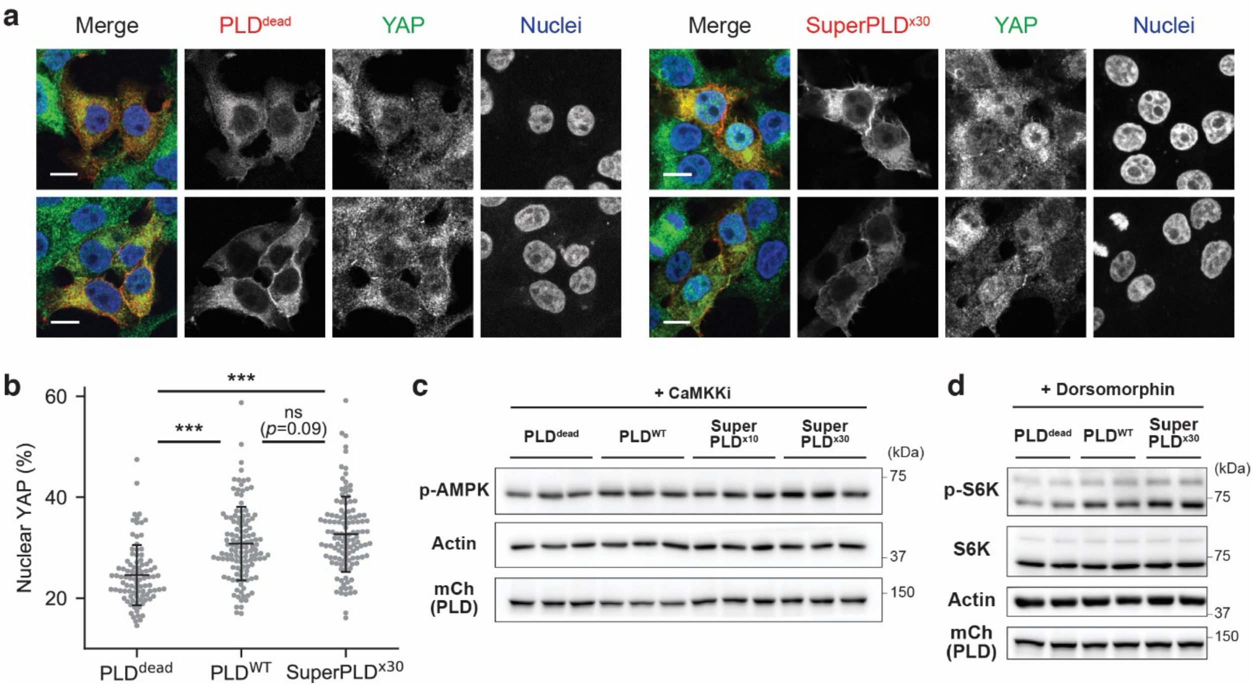

**Supplementary Figure 5. Application of superPLD to manipulate PA signaling. a–b**, Quantification of nuclear YAP level to evaluate Hippo signaling activity. HEK 293T cells expressing plasma membrane-targeted optoPLD (PLD$^{\text{dead}}$, PLD$^{\text{WT}}$ or superPLD$^{\text{x30}}$) were immunostained for YAP (**a**), and percent signal of YAP colocalized with DAPI (nucleus) signal is plotted for each transfected cell. Horizontal lines indicate average and vertical error bars indicate standard deviation (n=100–120). Scale bars: 10 µm. **c**, Representative Western blots used to quantify p-AMPK levels (see Fig. 5d). Cells expressing plasma membrane-targeted optoPLD were treated with an CaMKK inhibitor (STO-609) for 6 h to block CaMKK-mediated AMPK activation, followed by a 30 min incubation with 488 nm light. **d**, Representative Western blots used to quantify p-S6K levels (see Fig. 5e). Cells expressing plasma membrane-targeted optoPLD were treated with an AMPK inhibitor (dorsomorphin) for 1 h, followed by a 30 min incubation with 488 light.

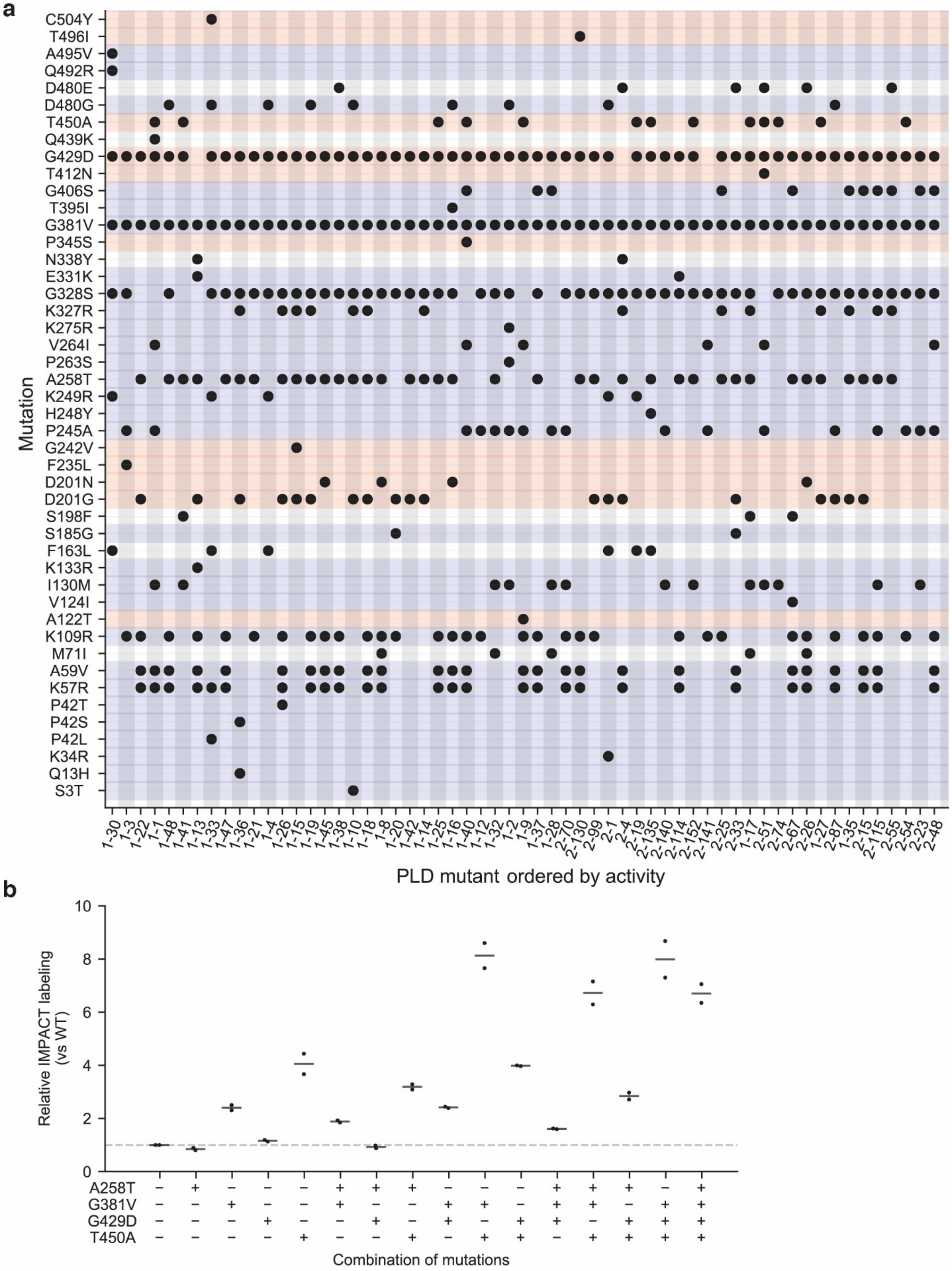
a
Mutation
C504Y
T496I
A495V
Q492R
D480E
D480G
T450A
Q439K
G429D
T412N
G406S
T395I
G381V
P345S
N338Y
E331K
G328S
K327R
K275R
V264I
P263S
A258T
K249R
H248Y
P245A
G242V
F235L
D201N
D201G
S198F
S185G
F163L
K133R
I130M
V124I
A122T
K109R
M71I
A59V
K57R
P42T
P42S
P42L
K34R
Q13H
S3T
PLD mutant ordered by activity
b
Relative IMPACT labeling (vs WT)
A258T
G381V
G429D
T450A
Combination of mutations

**Supplementary Figure 6. Mutations identified in various superPLD clones. a,** PLD mutant clones are shown in order of PLD activity in cells determined by IMPACT (increasing from left to right), and black dots indicate the presence of a particular point mutation in that PLD mutant. Mutants with identical sets of mutations are not shown in the plot. The mutated residues are colored based on conservation across PLDs from different species, determined by ConSurf[1–3]; red: high conservation, white: medium, blue: low. **b**, Activity assay of PLD mutants containing different combinations of four commonly occurring mutations that were generated in the $PLD^{WT}$ background (A258T, G381V, G429D and T450A). Cells expressing plasma membrane-targeted optoPLD with indicated set of mutations were labeled with IMPACT using 1 mM azidopropanol and 1 μM BCN-BODIPY. IMPACT fluorescence intensity normalized to optoPLD expression was determined by flow cytometry, and the relative values for each mutant compared to the $PLD^{WT}$ are plotted as relative IMPACT labeling (n=2 distinct samples). The effect of each mutation occurred mostly in a combinatorial manner (i.e., most mutations exhibited multiplicative effects in either increasing (G381V, T450A) or slightly decreasing (A258T) the activity, though G429D slightly increased activity alone but had negligible effects when combined with other mutations).

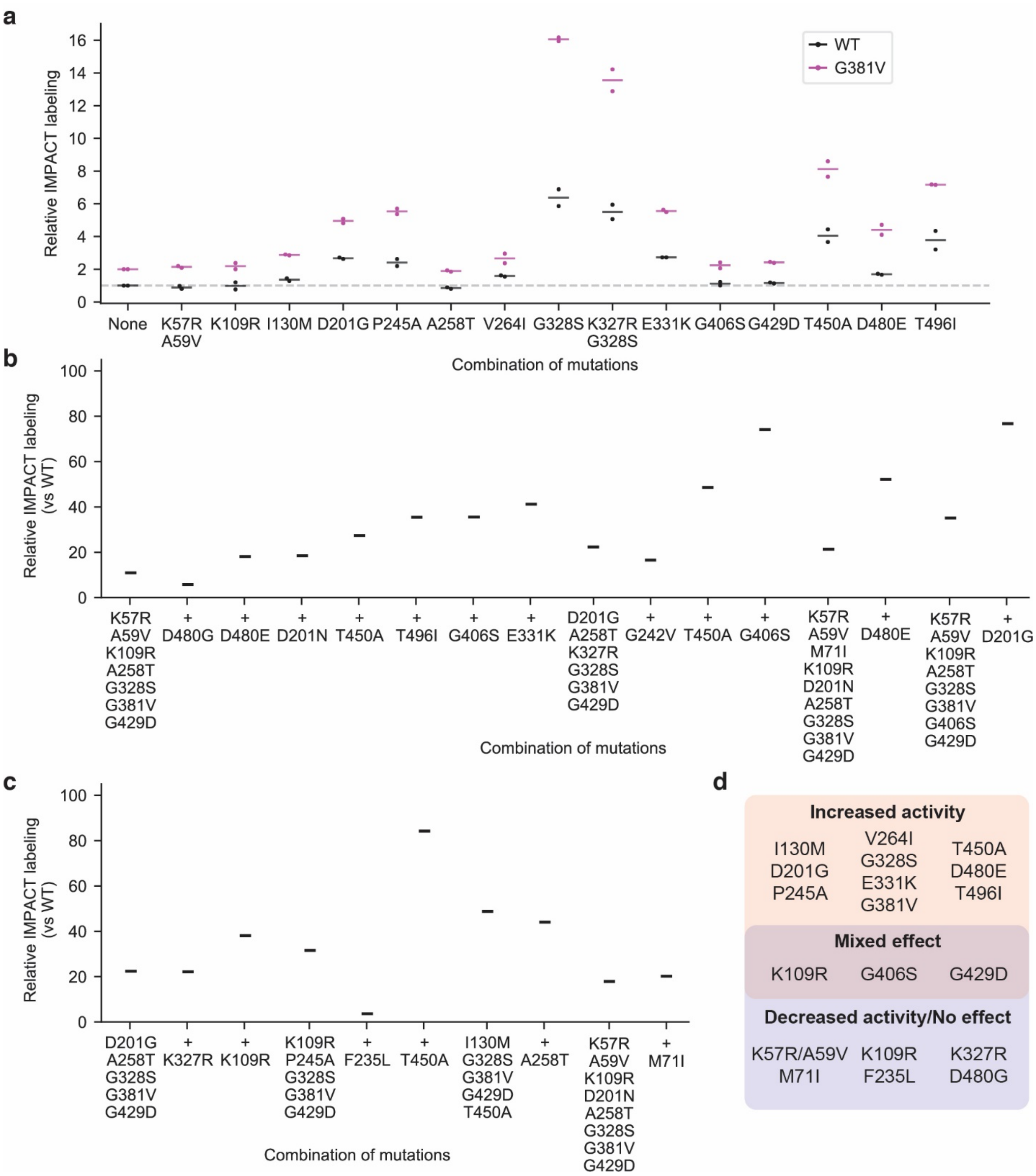

**Supplementary Figure 7. Analysis of the effects of acquired mutations on PLD activity. a**, Relative activity of PLDs with frequently occurring mutations that were individually installed into either $PLD^{WT}$ (black) or G381V (magenta) background. Cells were labeled and analyzed by flow cytometry as described in Supplementary Fig. 6. Horizontal lines indicate average (n=2) of relative mean intensities of IMPACT fluorescence. Dashed gray lines indicate the activity of $PLD^{WT}$, which is normalized to 1. **b–c**, Systematic comparison to evaluate the effect of individual mutations in the context of actual superPLD mutants isolated from the screen. The graphs show related mutants grouped separately. Each group contains a "template mutant" consisting of a specific set of mutations (which came from directed evolution experiments), which is shown left-most within each group, as well as other mutants that contain all the mutations in the template mutant plus one additional mutation that is indicated. **d**, Three different types of effects caused by individual mutations, determined based on point mutation analysis and systematic comparison analysis of mutants.

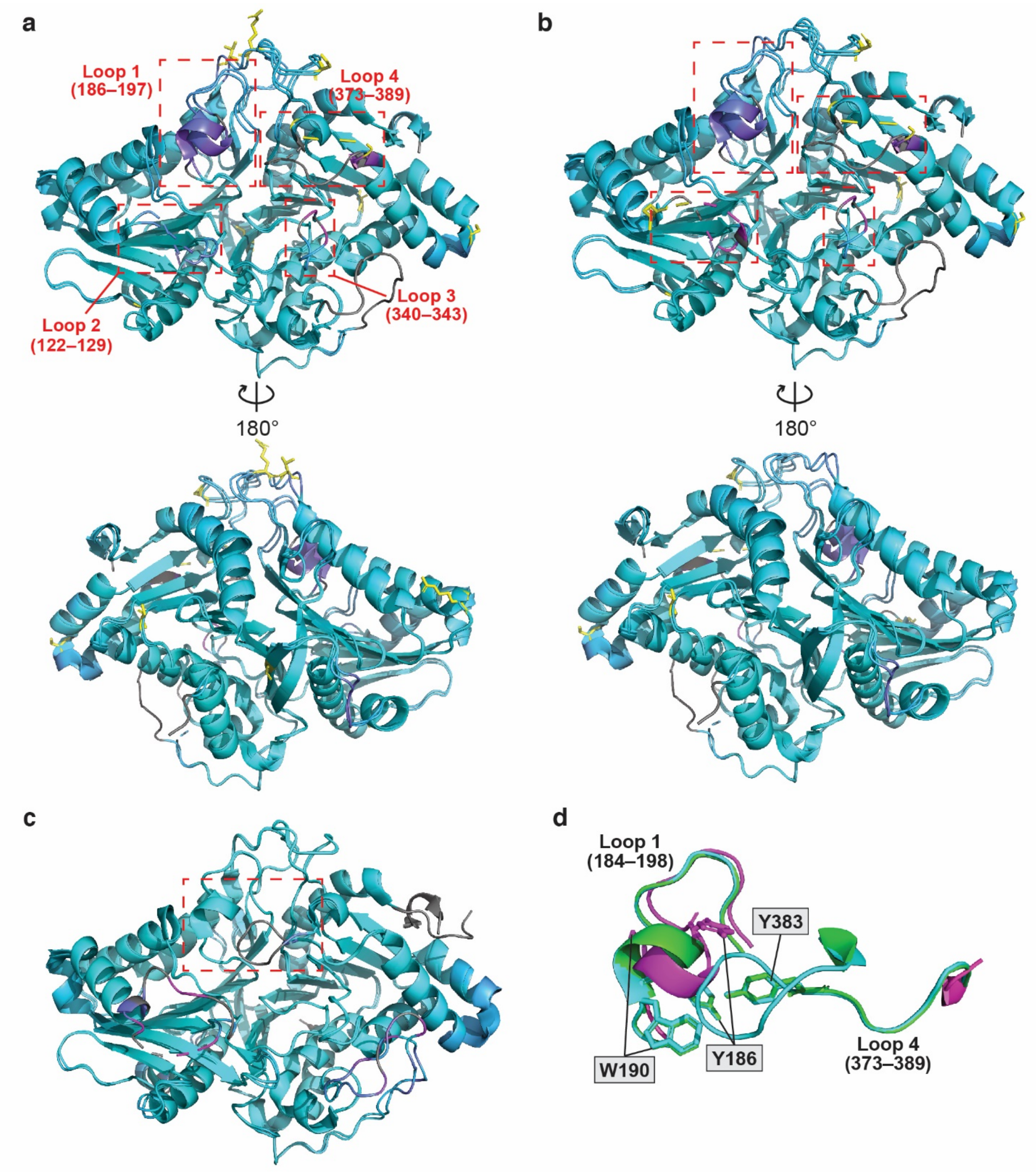

**Supplementary Figure 8. Mapping correlations between structural shifts and mutated sites in superPLDs. a–b**, Alignments of $PLD^{WT}$ with either superPLD (2-48) (**a**) or superPLD (2-23) (**b**), with the changes in two structures colored by root mean square deviation (RMSD) using the ColorByRMSD script[4]. The distances between aligned Cα atom pairs are stored as B-factors of these residues, which are colored by a color spectrum, with cyan specifying the minimum pairwise RMSD and magenta indicating the maximum. Unaligned residues are colored gray. Sites of mutations in each superPLD are shown in yellow, and mutated residues are shown as yellow sticks except for G381V, which exists on the missing flexible loop that is shown as a yellow dashed line.

Loops 1–4 at the entry of the active site are highlighted in red dashed boxes. **c**, Alignment of PLD from *Streptomyces sp. PMF* and PLD from *Streptomyces antibioticus*, with the differences between the two structures colored by RMSD, demonstrating a high level of structural homology. Loops 1 and 4 are highlighted in red dashed boxes. **d**, Zoomed-in image of loops 1 and 4. PLD from *Streptomyces sp. PMF*, PLD from *Streptomyces antibioticus*, and superPLD (2-48) are shown in green, cyan, and magenta, respectively. Note that the positions of residues Y186, W190 and Y383 are identical in two wildtype PLDs and found to be largely shifted in superPLD (except for Y383, which is unresolved but likely substantially displaced as discussed in the main text).

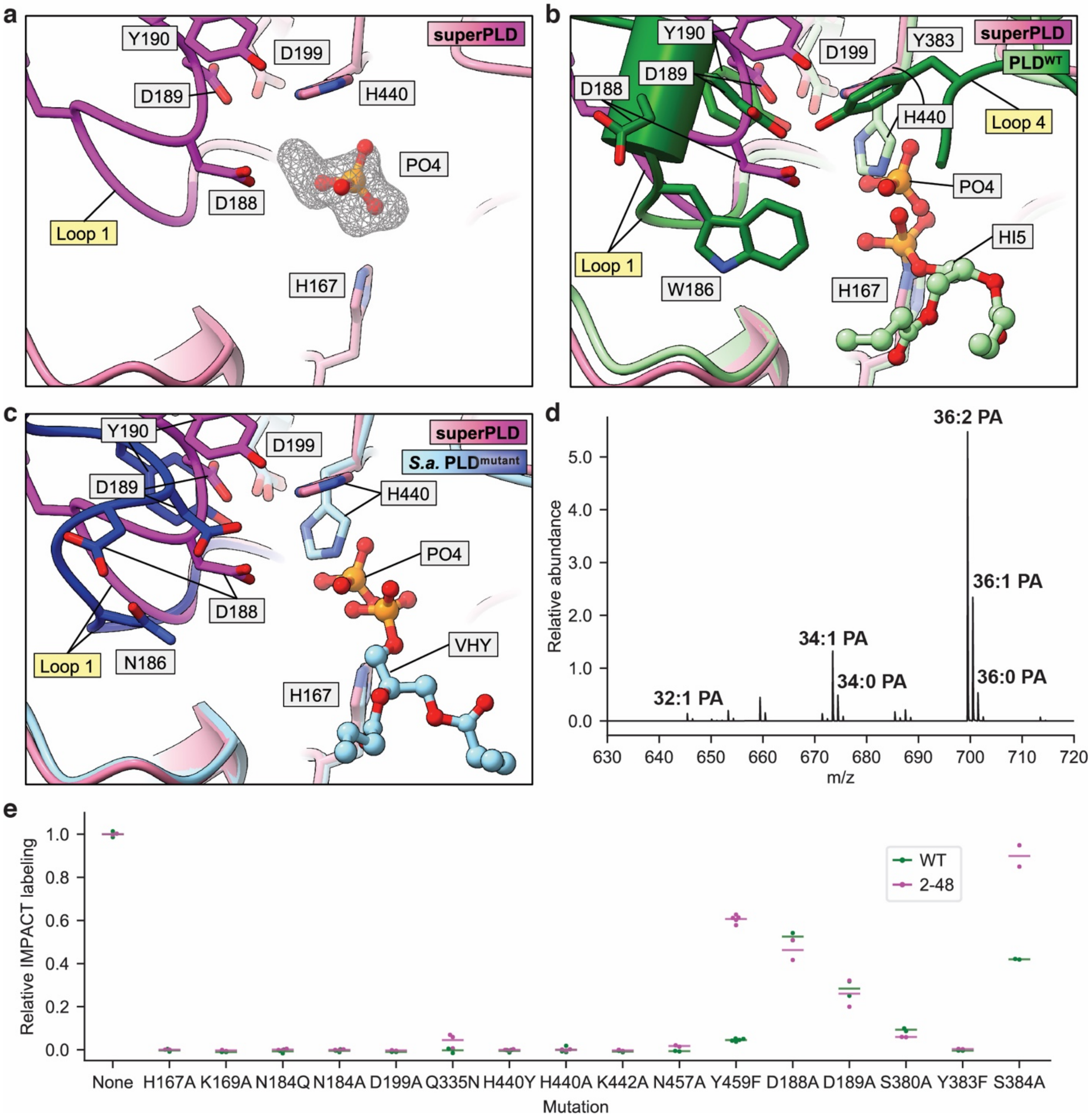

**Supplementary Figure 9. Analysis of the superPLD active site structure of superPLD. a–c**, Structures of the active sites of superPLD (2-48) (**a**), superPLD overlaid with PLD$^{WT}$ (PDB ID: 1V0Y; green) (**b**), and superPLD overlaid with PLD from *Streptomyces antibioticus* engineered to produce phosphatidylinositol[5] (PDB ID: 7JRV; light blue) (**c**). Loops 1 and 4 in each structure are shown in dark purple (superPLD 2-48) and dark blue (engineered *S. antibioticus* PLD). A phosphate ligand is modeled in the electron density found in the active site of superPLD. Polder omit map for the region around the modeled phosphate molecule is shown as a mesh, contoured at 0.52 σ using a carve distance of 3 Å[6]. **d**, LC–MS analysis of the lipid extract from purified superPLD, demonstrating the existence of multiple PA species that co-purified with superPLD (2-48). **e**, Comparison of the effects of site-directed mutation on PLD$^{WT}$ (green) vs. superPLD (2-48; magenta). HEK 293T cells expressing plasma membrane-targeted optoPLD versions of PLD$^{WT}$ or superPLD (2-48) with the indicated point mutation were labeled with 10 mM (for $PLD^{WT}$) or 100 μM (for superPLD) azidopropanol, followed by click chemistry tagging with 1 μM BCN-BODIPY. IMPACT fluorescence intensity normalized to optoPLD expression was determined by flow cytometry, and the relative values for each mutant compared to the appropriate parental PLD (i.e., $PLD^{WT}$ or superPLD (2-48)) are plotted as relative IMPACT labeling. Plots are replicates (n=2–5 distinct samples) from flow cytometry analysis, and the line indicates the average.

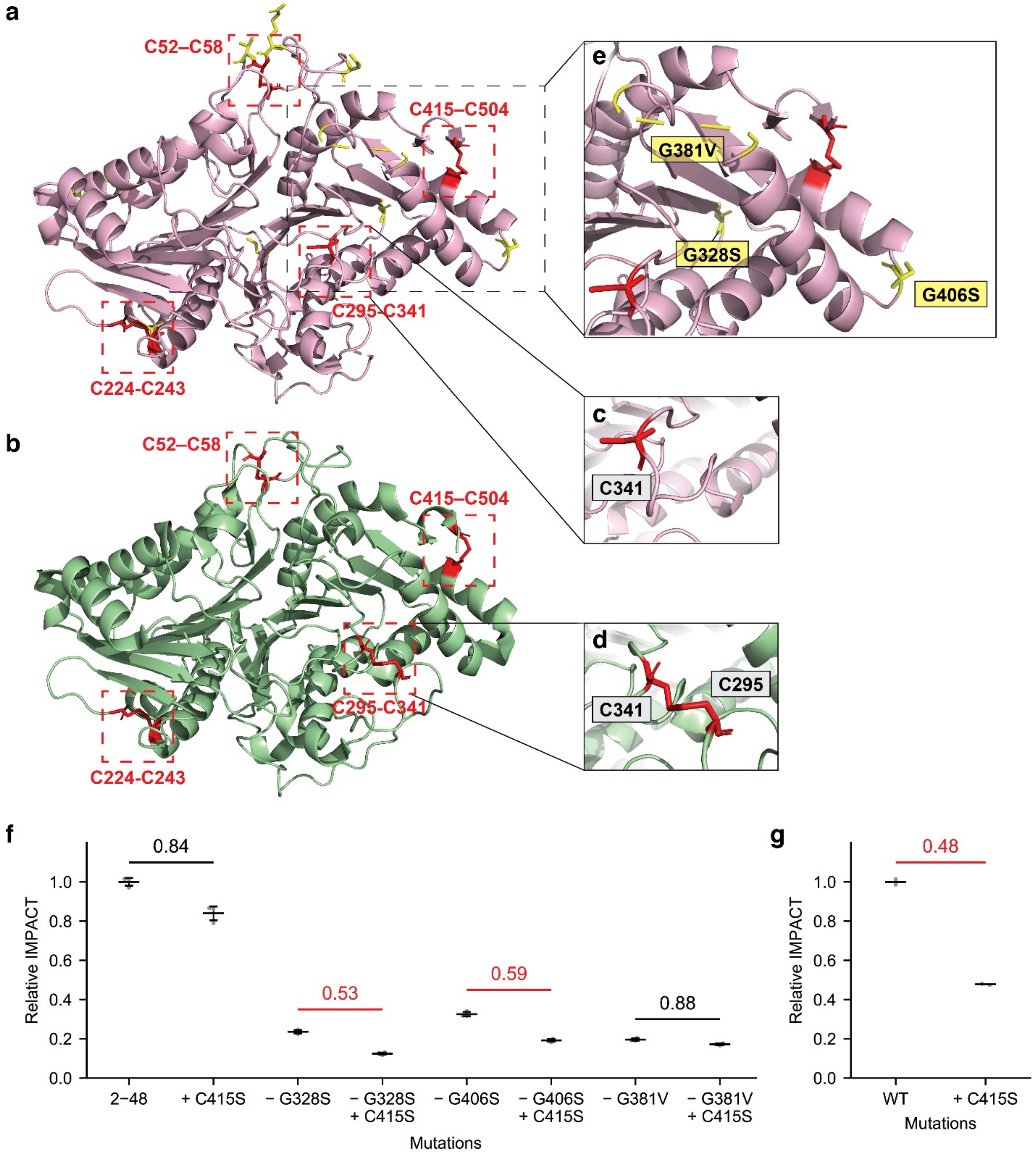

**Supplementary Figure 10. Disulfide bonds in superPLD and PLD$^{WT}$ structures. a–d**, Crystal structures of superPLD (2-48) (**a**, **c**, **e**) and PLD$^{WT}$ (PDB ID: 1V0Y) (**b**, **d**) with cysteine residues shown in red. Disulfide bonds in PLD$^{WT}$ are highlighted with red dashed boxes. In superPLD structures, C295 was not resolved, and C341 residue was flipped, indicating that the C295–C341 disulfide bond is reduced. Sites of mutations in superPLD are shown in yellow, with mutated residues shown as yellow sticks (except for G381V, which is in an unresolved flexible loop and is shown as a yellow dashed line). **f–g**, Identification of a mutation in superPLD that is responsible for enhanced tolerance to disabled disulfide bond formation. Mutations in superPLD (2-48) that occur at three positions near C415 (G328S, G381V, and G406S) were, within the superPLD (2-48) background, reverted to the residue that occurs in $PLD^{WT}$, and the relative effects of C415S mutation on PLD activity in these constructs (**f**) were compared to the effect of the C415S mutation in $PLD^{WT}$ (**g**). IMPACT fluorescence intensity normalized to optoPLD expression was determined by flow cytometry. Horizontal lines indicate average (n=3) of relative mean intensities of IMPACT fluorescence of cells expressing the indicated mutant PLD compared to the parental PLD (e.g., $PLD^{WT}$ or superPLD (2-48)), as measured by flow cytometry.

### Supplementary Tables

#### Supplementary Table 1. List of plasmids used in this study.

| | Name | Source | Notes |
|---|---|---|---|
| **1** | pCDNA-CRY2-mCherry-PLD-P2A-CIBN-CAAX | Addgene: 140114[7] | Used as a template for cloning |
| **2** | pCDNA-CRY2-mCherry-PLD$^{dead}$-P2A-CIBN-CAAX | Addgene: 140061[7] | Used as a template for cloning |
| **3** | GFP-PASS | Zhang et al., 2014[8] | Used as a template for cloning |
| **4** | pCDH-CRY2-mCherry-PLD-P2A-CIBN-CAAX | This study | Prepared by cloning CRY2-mCherry-PLD-P2A-CIBN-CAAX from **1** into a lentiviral vector with BamHI/EcoRI sites before/after PLD |
| **5** | pCDH-CRY2-mCherry-PLD$^{dead}$-P2A-CIBN-CAAX | This study | Prepared by cloning CRY2-mCherry-PLD$^{dead}$-P2A-CIBN-CAAX from **2** into a lentiviral vector with BamHI/EcoRI sites before/after PLD |
| **6** | pCDNA-CRY2-mCherry-B-PLD-E-P2A-CIBN-CAAX | This study | Prepared by introducing BamHI/EcoRI sites before/after PLD of **1** |
| 7 | pCDH-GFP-PASS | This study | Prepared by cloning GFP-PASS from **3** into a lentiviral vector |
| **8** | pCDH-CRY2-mCherry-PLD | This study | Prepared by cloning CRY2-mCherry-PLD from **4** into a lentiviral vector |
| **9** | pCDH-CRY2-mCherry-PLD$^{dead}$ | This study | Prepared by cloning CRY2-mCherry-PLD$^{dead}$ from **5** into a lentiviral vector |
| **10** | pCDH-CIBN-CAAX | This study | Prepared by cloning CIBN-CAAX from 1 into a lentiviral vector |
| **11** | pCDNA-p18-CIBN-P2A-CRY2-mCherry-PLD | This study | Prepared by cloning p18 (MGCCYSSENEDSDQDREERKLLLDPSSPPTKALNGAEPNY)[9], CIBN and CRY2-mCherry-PLD into pCDNA3.1 vector |
| **12** | pCDH-p18-CIBN | This study | Prepared by cloning p18-CIBN from **11** into a lentiviral vector |
| **13** | NusA-10xN-HRV3C (pCAV4.1) | Bosire et al., 2020[10] | Used as a vector for cloning |
| **14** | NusA-10xN-HRV3C-PLD | This study | Prepared by cloning PLD into **13** with BamHI/EcoRI sites before/after PLD |
| **15** | EGFP-LKB1 | This study | Prepared by cloning LKB1 into EGFP-C1 vector |

**Supplementary Table 2. List of primers for cloning plasmids used in this study.**

| | Name | Sequence | Notes |
|---|---|---|---|
| **1** | OL-CRY2-1-AS | GGAGAAAGTAGTGAAGTAGAGTTCCCAACAACTTT CTTG | For silencing existing EcoRI site in CRY2 in optoPLD |
| **2** | OL-CRY2-2-S | CAAGAAAGTTGTTGGGAACTCTACTTCACTACTTTC TCC | |
| **3** | OL-CRY2-2-AS | CGTCCCATGGATGATGAATCCATTCAGTTGGCAATC | |
| **4** | OL-CRY2-3-S | GATTGCCAACTGAATGGATTCATCATCCATGGGACG | |
| **5** | mCh-BamHI-PLD-AS | cagatccgccggatccCGATCGACCTGCTCCTCC | For introducing BamHI/EcoRI sites before/after PLD in optoPLD |
| **6** | mCh-BamHI-PLD-S | atcgggatccGGCGGATCTGCAGATTCAG | |
| **7** | PLD-EcoRI-P2A-AS | aagtttgtagcgccgcttccgaattcAGCGTTGCAGATTCCTCTTG | |
| **8** | BamHI-PLD-S | atcgggatccGGCGGATCTGCAGATTCAG | For amplification of PLD (Taq) |
| **9** | EcoRI-PLD-S | aagtttgtagcgccgcttccgaattcAGCGTTGCAGATTCCTCTTG | |
| **10** | BamHI-PLD-S | GCAGGTCGATCGGGATCC | For amplification of PLD (Phusion) |
| **11** | EcoRI-PLD-AS | GTAGCGCCGCTTCCGAATTC | |
| **12** | EcoRI-GFP-S | gtcaGAATTCGCCACCATGGTGAGCAAG | For cloning pCDH-GFP-PASS |
| **13** | BamHI-PASS-AS | GGTGGATCCTTAACTAGTCTTAGTGG | |
| **14** | pCDH-CRY2-S | tttgacctccatagaagattGCCACCATGAAGATGGAC | For cloning pCDH-CRY2-mCherry-PLD |
| **15** | pCDH-PLD-AS | cgcagatccttgcggccgcgCTAAGCGTTGCAGATTCCTC | |
| **16** | AgeI-p18-S | gtACCGGTATGGGGTGCTGCTACAGCAGCGAGAACG AGGACTCGGACCAGGACCGAGAGGAGCGGAAGCT GCTGCTGG | For swapping p18 with CAAX in optoPLD |
| **17** | ApaI-p18-AS | tGGGCCCTTAGTAGTTGGGCTCGGCTCCATTGAGAG CTTTGGTAGGGGGGCTGCTAGGGTCCAGCAGCAGC TTCCGCTCC | |
| **18** | vec-p18-S | tttaaacttaagcttggtaccgccaccATGGGGTGCTGCTACAGC | For cloning pCDNA3-p18-CIBN-P2A-CRY2-mCherry-PLD |
| **19** | p18-CIBN-AS | tcatggtaccGTAGTTGGGCTCGGCTCC | |
| **20** | p18-CIBN-S | gcccaactacGGTACCATGAATGGAGCTATAG | |
| **21** | CIBN-P2A-CRY2-AS | cgccagcctgCTTGAGCAGACTGAAGTTGGTAGCACCGC TTCCATGAATATAATCCGTTTTCTCC | |
| **22** | CIBN-P2A-CRY2-S | tctgctcaagCAGGCTGGCGATGTCGAGGAGAATCCAGG ACCTATGAAGATGGACAAAAAGAC | |
| **23** | PLD-taa-vec-AS | agtggatccgagctcggtacttaAGCGTTGCAGATTCCTC | |
| **24** | EcoRI-p18-S | GtcagaattcGCCACCATGGGGTGCTGC | For cloning pCDH-p18-CIBN |
| **25** | BamHI-CIBN-AS | gtcaggatccTCAATGAATATAATCCGTTTTCTCCAATTC | |
| **26** | pCAV4- PLD-S | aagttctgtttcagggtccgGGATCCGGCGGATCTGCA | For cloning NusA-10xN-HRV3C-PLD |
| **27** | pCAV4-PLD-AS | gagcctttcgttttatttgcttaGAATTCAGCGTTGCAGATTCCTC | |
| **28** | seq-mChterm(112)-S | CGTGGAACAGTACGAACGC | For sequencing PLD mutants |
| **29** | seq-CIBN(103)-AS | GGTCACCTCCTATAGCTCCATTC | |
| **30** | LKB1-S | gtcAGATCTATGGAGGTGGTGGACCCG | For cloning EGFP-LKB1 |
| **31** | LKB1-AS | gtGGTACCTCACTGCTGCTTGCAGGC | |

**Supplementary Table 3. List of primers for cloning PLD mutants used in this study.**

| | Name | Sequence |
|---|---|---|
| **1** | K109R-AS | CTTATTGCCTCTAGCTGCGGACTCCTTC |
| **2** | K109R-S | CCGCAGCTAGAGGCAATAAGTTGAAAGTTAGG |
| **3** | I130M-AS | CATGAATGTAATGCCTTCCAAGTATAGGGACG |
| **4** | I130M-S | ACTTGGAAGGCATTACATTCATGTGGTAGACGG |
| **5** | H167A-AS | GATTTTGCTGGCGTTCCAGGAG |
| **6** | H167A-S | CTGGAACGCCAGCAAAATCTTGG |
| **7** | K169A-AS | CCAAGATGGCGCTATGGTTCCAG |
| **8** | K169A-S | CCATAGCGCCATCTTGGTTGTAGAC |
| **9** | N184A-AS | CTTCCAGCTGGCTATTCCACCAGTAAG |
| **10** | N184A-S | GTGGAATAGCCAGCTGGAAGGATG |
| **11** | N184Q-AS | CTTCCAGCTCTGTATTCCACCAGTAAG |
| **12** | N184Q-S | GTGGAATACAGAGCTGGAAGGATG |
| **13** | D188A-AS | AAGTAATCGGCCTTCCAGCTATTTATTCC |
| **14** | D188A-S | GCTGGAAGGCCGATTACTTGGATAC |
| **15** | D189A-AS | ATCCAAGTAGGCATCCTTCCAGC |
| **16** | D189A-S | GGAAGGATGCCTACTTGGATACTACTC |
| **17** | D199A-AS | GATCAACGGCAGACACTGGATGAG |
| **18** | D199A-S | CAGTGTCTGCCGTTGATCTTGCAC |
| **19** | D201G-AS | TCTGATGTTGGTCTTGCACTGACCGGACC |
| **20** | D201G-S | CAGTGCAAGACCAACATCAGACACTGGATGAG |
| **21** | C224S-AS | CTTGTTTTGGGAGGTCCATGTCCAC |
| **22** | C224S-S | CATGGACCTCCCAAAACAAGAGTAAC |
| **23** | P245A-AS | GCATAGTGGCCATGCACCCTGCGTTG |
| **24** | P245A-S | GGTGCATGGCCACTATGCATAAAGATACAAACC |
| **25** | A258T-AS | CCCGGTGGTAGGTGATGCCTTGGGG |
| **26** | A258T-S | CATCACCTACCACCGGGAATGTCCC |
| **27** | V264I-AS | GAATGTCCCAATAATAGCCGTTGGAGGGT |
| **28** | V264I-S | ACGGCTATTATTGGGACATTCCCGGTG |
| **29** | K327R/G328S-AS | CAATGTGACTCCTTGCTGAAGCAACAAGAGC |
| **30** | K327R/G328S-S | CTTCAGCAAGGAGTCACATTGAAATATCTCAGCAG |
| **31** | G328S/E331K-AS | CCTGCTGAGATATTTTAATGTGACTTTTTGCTGAAGC |
| **32** | G328S-AS | CAATGTGACTTTTTGCTGAAGCAACAAGAG |
| **33** | G328S-S | CTTCAGCAAAAAGTCACATTGAAATATCTCAGCAG |
| **34** | E331K-AS | CCTGCTGAGATATTTTAATGTGACCTTTTGCTGAAGC |
| **35** | E331K-S | GTCACATTAAAATATCTCAGCAGGATTTGAAC |
| **36** | Q335N-AS | CGTTCAAATCGTTCTGAGATATTTCAATG |
| **37** | Q335N-S | GAAATATCTCAGAACGATTTGAACGCTAC |
| **38** | C341S-AS | AGGGTGGAGATGTAGCGTTC |

| | | |
|---|---|---|
| **39** | C341S-S | CGCTACATCTCCACCCTTGCC |
| **40** | S380A-AS | CCCCCAGCGCCCACTGC |
| **41** | S380A-G381V-AS | ACCCACAGCGCCCACTGC |
| **42** | S380A-G381V-S | GTGGGCGCTGTGGGTTACAG |
| **43** | S380A-S | GTGGGCGCTGGGGGTTACA |
| **44** | V381G-AS | GTAACCCCCACTGCCCACTGCGC |
| **45** | V381G-S | GGGCAGTGGGGGTTACAGCCAAATAAAATC |
| **46** | G381V-S384A-AS | TATTTGGGCGTAACCCACACTGC |
| **47** | G381V-S384A-S | GTGGGTTACGCCCAAATAAAATCACTTA |
| **48** | G381V-Y383F-AS | TTGGCTGGCACCCACACTGC |
| **49** | G381V-Y383F-S | GTGGGTGCCAGCCAAATAAAATCAC |
| **50** | Y383F-AS | TGGCTGGCACCCCCACTGC |
| **51** | Y383F-S | GGGGGTGCCAGCCAAATAAAATC |
| **52** | S384A-AS | TTTGGGCGTAACCCCCACTGC |
| **53** | S384A-S | GGGGGTTACGCCCAAATAAAATCAC |
| **54** | G406S-AS | CCCATTTGTCGTTGGGGGAGGATCTAAAG |
| **55** | G406S-S | CCCCAACGACAAATGGGCAGACGGGC |
| **56** | C415S-AS | GCAAATTGCTAGACATTGCAGTTTTG |
| **57** | C415S-S | CTGCAATGTCTAGCAATTTGCAGC |
| **58** | G429D-AS | CCCATTTGTCGTTGGGGGAGGATCTAAAG |
| **59** | G429D-S | CCCCAACGACAAATGGGCAGACGGGC |
| **60** | H440A-AS | CAACTTATGGGCTTGGGCATAAGGATG |
| **61** | H440A-S | CTTATGCCCAAGCCCATAAGTTGGTTTC |
| **62** | H440W-AS | CAACTTATGCCATTGGGCATAAGGATG |
| **63** | H440W-S | CTTATGCCCAATGGCATAAGTTGGTTTC |
| **64** | H440Y-AS | CAACTTATGGTATTGGGCATAAGGATG |
| **65** | H440Y-S | CTTATGCCCAATACCATAAGTTGGTTTC |
| **66** | K442A-AS | GAAACCAAGGCATGGTGTTGGG |
| **67** | K442A-S | AACACCATGCCTTGGTTTCCGTC |
| **68** | T450A-AS | CGATATAAAATGCGGATGAGTCGACGGAAAC |
| **69** | T450A-S | GACTCATCCGCATTTTATATCGGTTCAAAAAATTTGTAC |
| **70** | N457A-AS | GATGGGTACAAGGCTTTTGAACCG |
| **71** | N457A-S | GTTCAAAAGCCTTGTACCCATCCTG |
| **72** | Y459F-AS | CCAGGATGGGAACAAATTTTTTGAAC |
| **73** | Y459F-S | CAAAAAATTTGTTCCCATCCTGGCTG |
| **74** | D480E-AS | CAATTTGGCCTCCAGTTGCTTGGCAGC |
| **75** | D480E-S | GCAACTGGAGGCCAAATTGCTTGACCCTC |
| **76** | D480G-AS | AGCAACTGGGTGCCAAATTGCTTGACCCTC |
| **77** | D480G-S | CAATTTGGCACCCAGTTGCTTGGCAGC |
| **78** | T496I-AS | GTAATCCACTATTGCAGTCTCCTGTGAGTAC |
| **79** | T496I-S | GAGACTGCAATAGTGGATTACGCAAGAGGAAT |

| | | |
|---|---|---|
| **80** | C52S-AS | CGCCCCAAGATCCTGGGG |
| **81** | C52S-S | CCAGGATCTTGGGGCGATG |
| **82** | K57R/A59V-AS | GATGATAGATGCGTTGACAGAGTTGGGACAAAAAG |
| **83** | K57R/A59V-S | TCTGTCAACGCATCTATCATCGCCCCAACATCCTG |
| **84** | C504S-AS | gtcaGAATTCAGCGTTGGAGATTCCTCTTGCG |

**Supplementary Table 4. Statistics from X-ray data collection, processing, and refinement.**

| | SuperPLD (2-23) | SuperPLD (2-48) |
|---|---|---|
| **Data collection** | | |
| Space group | *P*21 21 21 | *P*21 21 21 |
| Cell dimensions | | |
| *a*, *b*, *c* (Å) | 72.725, 73, 87.482 | 72.828, 73.254, 87.19 |
| *α*, *β*, *γ* (°) | 90, 90, 90 | 90, 90, 90 |
| Wavelength (Å) | | |
| Resolution (Å) | 56.05-1.905 (1.973-1.905) | 56.09-1.852 (1.918-1.852) |
| $R_{meas}$ (%) | 11.9 (69.6) | 12.25 (95.86) |
| I/σ (I) | 12.1 (2.0) | 13.55 (2.35) |
| Completeness (%) | 98.8 (94.9) | 99.15 (91.44) |
| Redundancy | 5.5 (5.4) | 13.1 (12.4) |
| $CC_{1/2}$ | 0.992 (0.820) | 0.998 (0.879) |
| **Refinement** | | |
| Resolution (Å) | 56.05-1.905 | 56.09-1.852 |
| No. of reflections | 36421 (3442) | 40025 (3612) |
| Completeness (%) | 98.8 | 99.15 |
| $R_{work}$ / $R_{free}$ (%) | 17.72/23.18 | 17.29/21.68 |
| **No. of non-hydrogen atoms** | 3842 | 3920 |
| Protein | 3624 | 3631 |
| Ligand | 5 | 5 |
| Water | 213 | 284 |
| **Wilson B-value (Å$^2$)** | 34.1 | 28.98 |
| **Mean B-value (Å$^2$)** | 44.28 | 36.59 |
| Protein | 44.21 | 36.29 |
| Ligand | 88.94 | 84.09 |
| Water | 44.4 | 39.58 |
| **Molprobity clash score** | 4.46 | 5.42 |
| **Ramachandran** Favored (%) | 96.2 | 95.78 |
| Allowed (%) | 2.95 | 3.38 |
| Outliers (%) | 0.84 | 0.84 |
| **RMS deviations** Bonds (Å) | 0.011 | 0.01 |
| Angles (°) | 1.12 | 1.08 |
